# Beneficial and detrimental consequences of AHR activation in intestinal infection

**DOI:** 10.1101/2025.05.28.656570

**Authors:** Oscar E. Diaz, Liang Zhou, Christopher Barrington, Dennis Lindqvist, Frederike Graelmann, Emma Wincent, Brigitta Stockinger

## Abstract

The ligand dependent transcription factor aryl hydrocarbon receptor (AHR) is an environmental sensor whose activation can have physiologically beneficial or detrimental consequences for host immune responses depending on the ligand. Here we investigated the hypothesis that prolonged AHR activation either due to inefficient ligand metabolism or due to genetic manipulation may underlie the distinction between beneficial and detrimental effects. Our data indicate that prolonged AHR activation caused toxic endpoints for liver and thymus but was not per se interfering with the host response to infection with the intestinal pathogen *C.rodentium*. Genetically driven constitutive AHR activation improved resistance to infection, whereas prolonged AHR activation by the pollutant TCDD resulted in delayed clearance of *C.rodentium* associated with a suppression in antibody production. Combined single cell RNAseq and ATAC-seq analysis provided evidence that TCDD, but not genetic AHR activation, negatively affected dendritic cell functions such as activation, maturation and antigen presentation. Thus, the detrimental impact of environmental pollutants such as TCDD on immune responses cannot solely be attributed to aberrantly prolonged activation of AHR.

## Introduction

The aryl hydrocarbon receptor (AHR) is a ligand dependent transcription factor that functions as a sensor of environmental pollutants but also recognises dietary and microbial ligands. It is evolutionary conserved in most bilaterian animals with important roles in development of sensory neurons in invertebrates such as *Drosophila* and *C. elegans* (Hahn, 2002; Hahn *et al*, 2017). In vertebrates research on AHR was initially focused on its role in the toxicity of man-made pollutants such as polychlorinated biphenyls (PCBs), dioxins, and polycyclic aromatic hydrocarbons (PAHs), and their detoxification by induction of biotransformation enzymes such as the cytochrome P450 family 1 (Cyp1). The toxicities associated with exposure to dioxins such as 2,3,7,8-tetrachlorodibenzo-*p-*dioxin (TCDD) and related compounds (Poland & Knutson, 1982), however, also pointed to important physiological functions in addition to inducing biotransformation enzymes.

It is now clear that the AHR has important functions, for instance, in mucosal sites such as the lung and gastrointestinal tract which are continuously exposed to ligands derived from external sources such as pollutants or diet or endogenous ligands generated from tryptophan metabolism by the constituents of the microbiota (Stockinger *et al*, 2021). Nevertheless, there is no clear consensus why and how environmental pollutants affect physiological AHR functions.

Several studies have addressed the detrimental effects of AHR activation by TCDD during influenza infection of the lung, such as suppression of innate and adaptive immune responses (Boule *et al*, 2018; Franchini *et al*, 2019; Houser *et al*, 2024; Houser & Lawrence, 2022). In contrast, exposure to the endogenous tryptophan metabolite formylindolo[3,2-b]carbazole (FICZ) which has similar affinity for AHR did not have such effects on the viral immune response (Wheeler *et al*, 2014). One of the prevailing theories explaining such differences is that the detrimental effects of TCDD is due to abnormally prolonged AHR stimulation (Bock, 2019; Mitchell & Elferink, 2009). While the endogenous ligand FICZ is rapidly metabolised by AHR induced Cyp1 enzymes (Bergander *et al*, 2004; Wincent *et al*, 2009) and therefore only activates AHR during a short time window, TCDD is not a good substrate for these enzymes and causes AHR activation persisting over weeks in mice (Birnbaum, 1986).

In this study we investigated whether prolonged AHR activation is indeed the root cause for detrimental effects in host responses to infection. For this we used two distinct models of prolonged AHR activation: a genetic model of constitutive AHR activation (Ye *et al*, 2017) as well as the AHR ligand TCDD. The experimental model we studied was infection with the pathogen *Citrobacter rodentium* as our previous data had shown the importance of AHR in recovery from this infection (Metidji *et al*, 2018; Schiering *et al*, 2017).

Contrary to our expectations that prolonged AHR activation is detrimental to bacterial infections, mice with constitutive AHR activation were able to clear *C.rodentium* faster than wildtype mice, whereas mice exposed to TCDD during infection showed delayed clearance and strongly reduced antibody responses to *C. rodentium* compared with wildtype mice. While previous studies showed that TCDD likely affects B cells directly (De Abrew *et al*, 2011), we suspected that TCDD might affect also affect a cell type that is required for induction of adaptive immune responses of both T and B cells, such as dendritic cells, as TCDD may disrupt dendritic cell homeostasis and function with consequences for the induction of T effector cells (Bankoti *et al*, 2010; Franchini *et al*., 2019). Using single cell profiling of gene expression and chromatin accessibility of immune cells from colon and colon draining mesenteric lymph node (c-MLN), our data show that TCDD, but not constitutively active AHR adversely affected the function of dendritic cells, in particular antigen presentation capacity and migration.

Thus, the duration of AHR signaling per se is not the decisive factor that characterises the detrimental effect of AHR activation by TCDD. However, both scenarios of prolonged AHR activation produced the known toxic effects on liver and thymus, highlighting divergence of toxicity and effects on immune reactivity.

## Results

### Differential AHR expression on gut cell populations

AHR is widely expressed in the immune system in a cell type and context specific manner. To visualise AHR protein expression on a single cell level by flow cytometry, we employed mice expressing a Td-Tomato fluorochrome knocked into the AHR locus (Diny et al, 2022). Fig.1 shows a comparison of AHR levels on different immune cell types in colon and c-MLN. AHR expression levels were generally higher in cells from the colon and the highest expression was seen in the myeloid lineage comprising dendritic cells, macrophages and eosinophils, confirming what was reported previously in the small intestine (Appendix Fig.S1) (Diny *et al*., 2022).

**Figure 1-.**
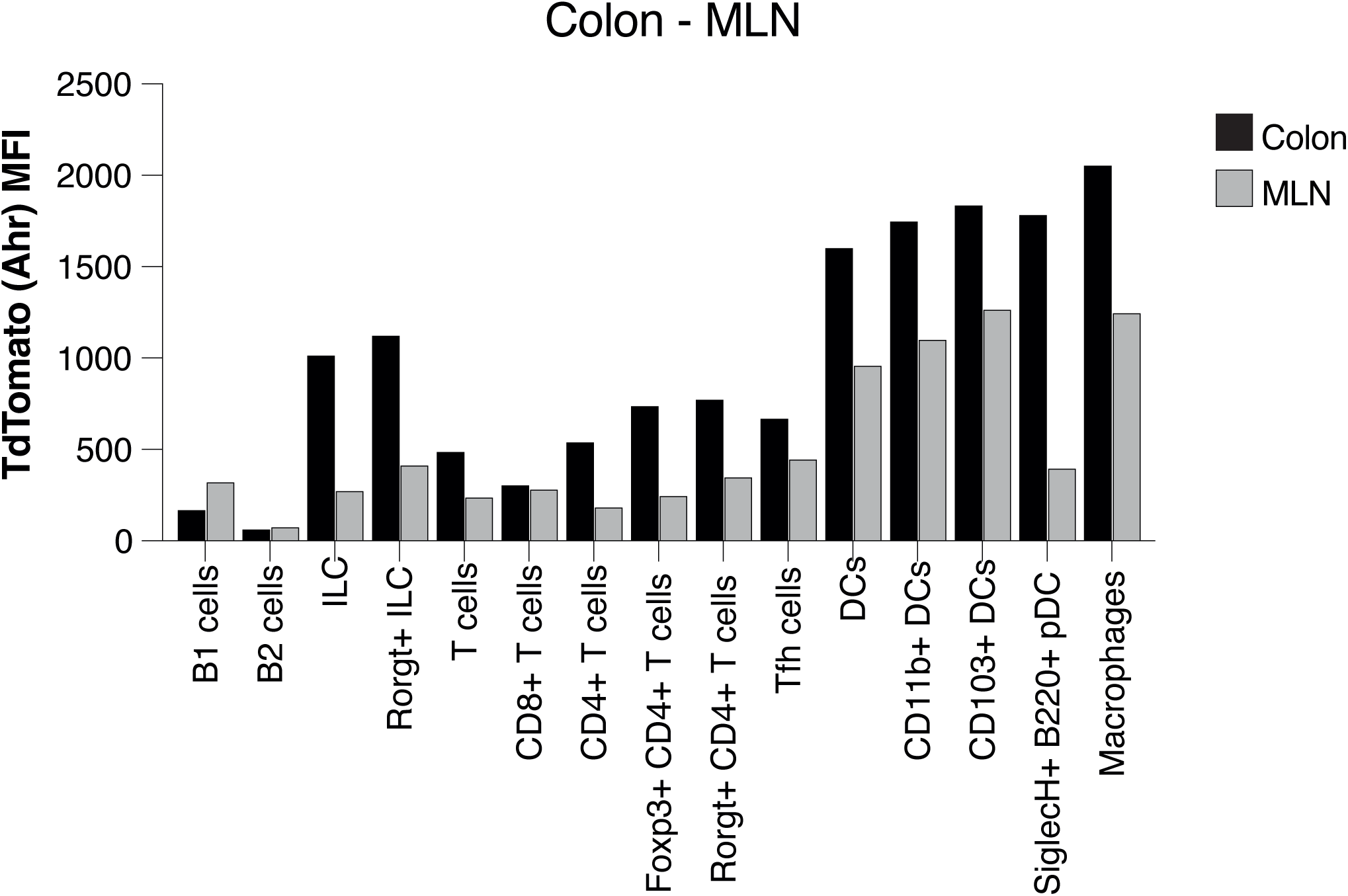
AHR expression in intestinal immune cell populations. AHR-TdTomato expression across immune cell types in the colon lamina propria and colon-draining lymph node was determined by flow cytometry. Bar plots represent the geometric MFI of concatenated events from cells obtained from n=2 mice per group. Representative of 3 experiments.

### Prolonged AHR activation in *Ahr^dCAIR/dCAIR^* and TCDD treated mice

Next we analysed the state of AHR activity in the genetic and TCDD model. In contrast to previous models of constitutively active AHR (CA-Ahr) which used a transgenic approach under the control of artificial genes and promoters outside of the Ahr locus (Andersson *et al*, 2002), the mouse model used here was generated using a knockin strategy into the endogenous locus. Specifically, a Flag-CA-Ahr-IRES-GFP (CAIR) allele preceded by a lox-stop-lox was introduced into the endogenous Ahr locus. Crossing with PGK-Cre led to the deletion of the STOP cassette so that mice expressed constitutively active AHR (dCAIR) in all tissues (Ye *et al*., 2017), hereafter referred to as *Ahr^dCAIR/dCAIR^*. Homozygous *Ahr^dCAIR/dCAIR^* mice were used in all experiments and compared with wildtype (WT) B6 mice that received a single dose of 10μg/kg TCDD by oral gavage.

Fig.2A shows constitutive expression of the AHR target gene *Cyp1a1* (Hu *et al*, 2007) in the colon of untreated *Ahr^dCAIR/dCAIR^* mice, indicative of constant AHR activation. *Cyp1a1* expression in WT mice that had received TCDD 6 days before analysis was of comparable magnitude (Fig.2B) along the gastrointestinal tract whereas vehicle treated mice displayed a diminishing gradient of *Cyp1a1* expression from proximal small intestine to colon (Fig.2C).

**Figure 2-.**
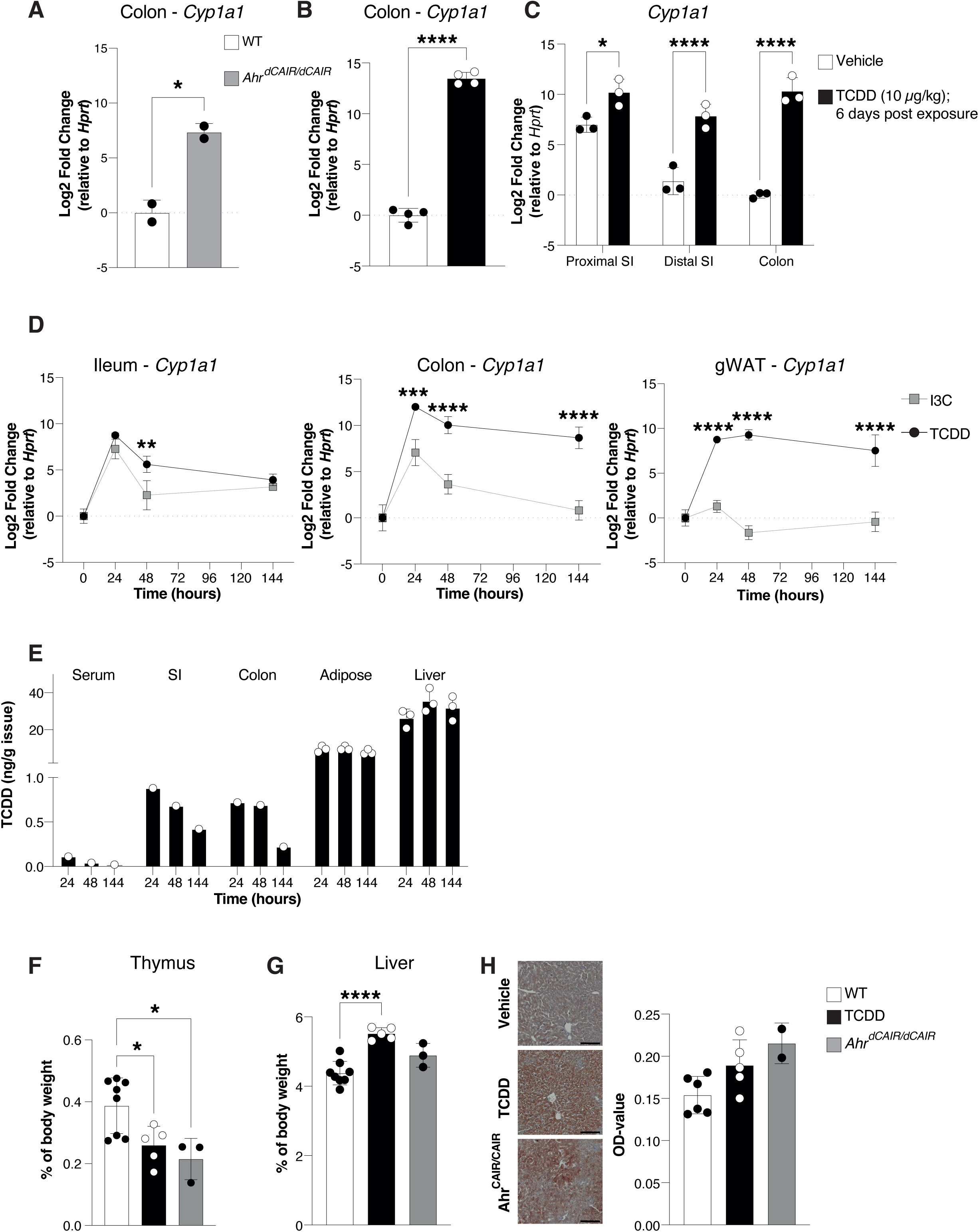
Prolonged AHR activation. (**A-C**) *Cyp1a1* gene expression was determined by qPCR from the indicated intestinal sections from *Ahr^dCAIR/dCAIR^* mice and WT mice (**A**) and 6 days after a single administration of 10 μg/kg TCDD (**B-C**) and presented as log2 Fold change relative to *Hprt*. Each dot represents one mouse and the bars show the mean + SD. Data correspond to 1 experiment with n=2-4 per group. Representative of 2 experiments. (**D**) *Cyp1a1* gene expression from the indicated organs was determined by qPCR. Timepoints correspond to the hours after a single administration of 250 mg/kg I3C or 10 μg/kg TCDD in corn oil. Expression values were normalized to *Hprt* and to values from the vehicle-treated mice. Each dot represents the mean of 2-3 mice + SD. (**E**) TCDD levels detected in the indicated organ and timepoint following a single administration of 10 μg/kg TCDD in corn oil normalized to the organ weight. Each dot represents one mouse for adipose and liver samples (n=3 per group) and a pool of samples from 3 mice for the serum, small intestine (duodenum and ileum) and colon. (**F-G**) Weight of thymus and liver of thymus and liver from *Ahr^dCAIR/dCAIR^*or TCDD treated mice. Each dot represents one mouse and the bars show the mean + SD. Data are from 1 experiment with n=3-5 per group. (**H**) Representative Oil Red O (ORO) staining in liver sections of *Ahr^dCAIR/dCAIR^* mice and TCDD-treated mice. Scale bars, 50 μm (left panel). Quantification of ORO stain (right panel). Each dot represents one mouse and the bars show the mean + SD. Data are from 1 experiment with n=3-5 per group. *p<0.05, **p<0.01, ***p<0.001, ****p<0.0001. Student’s t-test (A, B), Two-way ANOVA with Šídák’s multiple comparisons test (C, D), One-way ANOVA with Tukey’s multiple comparisons test (F-H).

We next compared the kinetics of *Cyp1a1* expression induced by TCDD in ileum, colon and gonadal white adipose tissue (gWAT) over a period of 6 days with that following oral application of the proligand indole 3 carbinol (I3C), a dietary constituent that is converted to the high affinity AHR ligand indolo[3,2-b]carbazole (ICZ) in the acidic environment of the stomach (Bjeldanes *et al*, 1991). *Cyp1a1* levels induced by both TCDD and I3C peaked at 24h, but while I3C induced *Cyp1a1* rapidly declined to baseline levels, TCDD induced *Cyp1a1* expression remained constant, indicative of prolonged AHR signaling (Fig.2D). In adipose tissue *Cyp1a1* induction was only seen after TCDD application. TCDD levels themselves declined over 6 days in small intestine and colon tissue, but remained constant in adipose tissue and particularly accumulated in the liver as previously described (Gasiewicz *et al*, 1983) (Fig.2E). I3C or its AHR agonist derivate ICZ were not quantifiable in any of the analysed tissues and timepoints.

TCDD toxicity is associated with hepatomegaly, intrahepatic lipid accumulation and thymic involution (Poland & Knutson, 1982). In order to assess whether prolonged AHR activation per se is responsible for toxicity characteristic for TCDD, we compared *Ahr^dCAIR/dCAIR^*mice with mice given a single dose of TCDD. Fig. 2F-H clearly shows that both models of prolonged AHR signaling exhibited clear toxic endpoints such as thymic involution and intrahepatic lipid accumulation, whereas hepatomegaly was not evident in *Ahr^dCAIR/dCAIR^* mice.

Thus, with respect to these classical toxic endpoints the two models were comparable.

### Effect of prolonged AHR activation on host response to infection

To evaluate the impact of prolonged AHR activation on the host response we infected mice with the intestinal pathogen *C. rodentium* which is a widely used model organism to study pathology as well as the host response to infection. There is a wealth of information about the kinetics of infection and the host innate and adaptive immune responses required to clear the pathogen (Basu *et al*, 2012; Caballero-Flores *et al*, 2021; Zindl *et al*, 2022). Furthermore, the importance of AHR signaling has been well documented in this model (Metidji *et al*., 2018; Schiering *et al*., 2017).

Upon infection with *C. rodentium, Ahr^dCAIR/dCAIR^* mice cleared the bacteria faster than wildtype mice (Fig.3A), made more IL-22 (Fig.3B) and exhibited lower inflammation indicated by fecal Lipocalin-2 (Lcn2) expression (Fig.3C). Furthermore, these mice were much more resistant to epithelial damage caused by dextran sulfate sodium (DSS) application, showing barely any weight loss and strongly reduced Lipocalin-2 levels (Fig.EV1A-C).

**Figure 3-.**
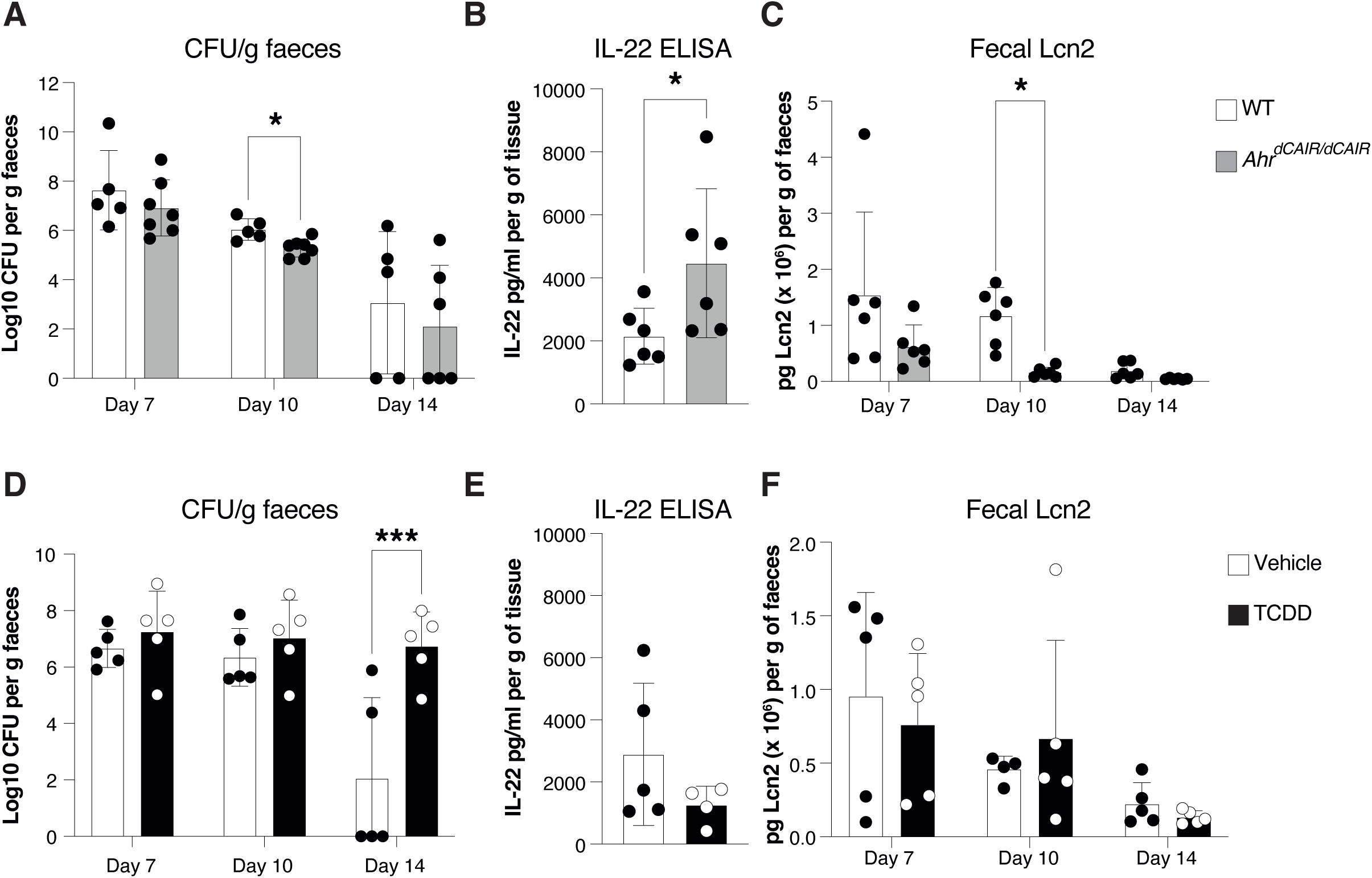
Impact of prolonged AHR activation to *Citrobacter rodentium* infection. (**A**) *C. rodentium* burden in faecal pellets, (**B**) IL-22 protein levels in supernatants from colon explant cultures, (**C**) Lcn2 protein levels in faecal pellets from *Ahr^dCAIR/dCAIR^* and WT mice (n=6-8 per group), representative of 2 experiments. (**D**) *C. rodentium* burden in faecal pellets, (**E**) IL-22 protein levels in supernatants from colon explant cultures and (**F**) Lcn2 protein levels in faecal pellets from TCDD-treated and vehicle-treated mice. Mice were infected 6 days after TCDD or vehicle administration, n=4-5 per group and representative of 2-3 experiments. Each dot represents samples collected from one mouse and the bars show the mean + SD. Two-way ANOVA with Šídák’s multiple comparisons test (A, C, D, F), Student’s t-test (B, E). *p<0.05, **p<0.01, ***p<0.001.

These data suggest that the prolonged activation of AHR in *Ahr^dCAIR/dCAIR^*mice did not negatively influence the host response against *C. rodentium* infection and instead accelerated the response without a concomitant increase in inflammation.

In contrast, mice treated with TCDD and infected with *C. rodentium* 6 days later exhibited delayed clearance of bacteria between day 7 and day 14, when most vehicle treated mice had eliminated the pathogen (Fig.3D). These data were obtained with male mice, but as females responded similarly (Fig.EV2A) males and females were pooled in subsequent experiments. Nevertheless, beyond day 21 *C. rodentium* was eventually also cleared in TCDD treated mice (Fig.EV2B). IL-22 levels were lower in TCDD treated mice (Fig.3E) and inflammation measured by Lipocalin-2 fecal expression was comparable to that seen in infected mice that had not received TCDD (Fig.3F).

These data indicate that TCDD exposure which similarly to *Ahr^dCAIR/dCAIR^*causes prolonged AHR activation had a detrimental effect on the host response to infection. In contrast, the negative influence of TCDD in *C. rodentium* infection was not replicated in the DSS colitis model where TCDD treated mice exhibited less weight loss and the overall levels of Lipocalin-2 declined faster than in mice that had not received TCDD (Fig.EV1D-F). Given that the DSS model is primarily a model for epithelial cell damage, this suggests that epithelial cells may not be the primary target for the adverse effects of TCDD (Chassaing *et al*, 2014; Yang & Merlin, 2024).

### Effect of prolonged AHR activation on humoral immunity

It is well established that a B cell response of IgG antibodies to *C. rodentium* is essential for effective clearance of the pathogen (Maaser *et al*, 2004) and TCDD was reported to suppress B cell responses (Boule *et al*., 2018; Houser & Lawrence, 2022; Sherr & Monti, 2013; Vorderstrasse *et al*, 2001). We therefore analysed the sera of mice exposed to TCDD 6 days before infection compared with infected vehicle treated and *Ahr^dCAIR/dCAIR^* mice.

Sera analysed on day 14 after infection showed that TCDD treated mice had significantly lower antibody responses to *C. rodentium* than controls mainly affecting the IgG2b, IgG3, IgM subclasses, whereas IgG1, IgG2c and IgA responses were not affected (Fig.4A, Appendix Fig.S2A).

**Figure 4-.**
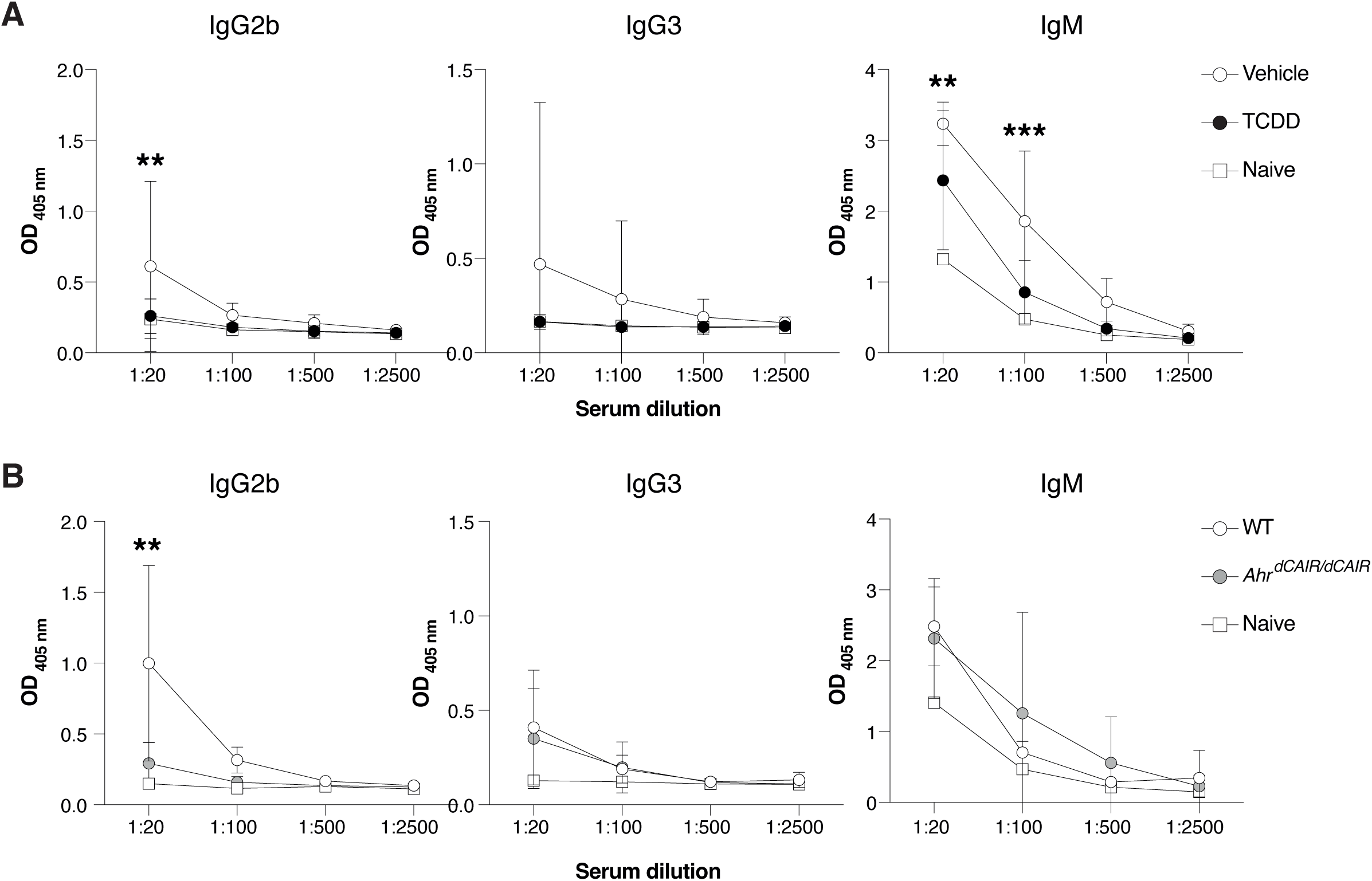
TCDD reduces levels of *C. rodentium*-specific antibodies. (**A**) Levels of *C.rodentium*-specific Ig subclasses at the indicated sera dilutions 14 days after *C. rodentium* infection in mice that received 10 μg/kg TCDD or vehicle 6 days before infection. Data are and representative of 2 experiments (vehicle, n=11; TCDD, n=10, Naïve =2). Each dot represents the mean + SD. (**B**) Levels of indicated Ig subclasses at the indicated sera dilutions 14 days after *C. rodentium* infection in *Ahr^dCAIR/dCAIR^* and WT mice. Data are from 2 experiments (WT, n=5; *Ahr^dCAIR/dCAIR^*, n=4, Naïve =1). Each dot represents the mean + SD. Two-way ANOVA with Tukey’s multiple comparisons test (A, B). *p<0.05, **p<0.01, ***p<0.001.

Conversely, antibody responses in *Ahr^dCAIR/dCAIR^* mice were comparable to control mice apart from the IgG2b response (Fig. 4B and Appendix Fig.S2B). While the antibody responses measured were quite variable, these results nevertheless suggest that TCDD has a broader suppressive effect than constitutive AHR activity in *Ahr^dCAIR/dCAIR^*mice. This was supported by similar reductions in T dependent and independent antibody responses to unrelated antigens such as NP-CGG and NP-Ficoll following TCDD application (Fig.EV3A,B).

### Single cell profiling highlights dendritic cell impairment by TCDD

Analysis of the c-MLN of TCDD treated mice 14 days after infection indicated a decrease in total CD45^+^ immune cells, particularly evident for dendritic cell populations and B cell subsets (Fig.EV4). These findings were reminiscent of previous reports in influenza infection of the lung following TCDD treatment (Houser *et al*., 2024; Vorderstrasse & Kerkvliet, 2001). We therefore decided to embark on profiling single cells isolated from colon and c-MLN on day 5 after infection with *C. rodentium,* a time when the adaptive immune response is activated. Our focus was on dendritic cells as the orchestrators for both arms of the adaptive immune response and in view of their exceptionally high AHR expression which makes them likely targets for environmental effects acting through this receptor.

As dendritic cells are present in very low numbers in these tissues, we deliberately enriched the populations submitted to single cell profiling for dendritic cells to increase their detection threshold. We performed simultaneous single nuclei ATAC and RNA-seq with a 1:1 mix of CD45^+^ cells and dendritic cells sorted from the colon and c-MLN of *Ahr^dCAIR/dCAIR^* and TCDD-exposed mice, as well as vehicle treated control mice at day 5 following *C. rodentium* infection (Fig.5A).

**Figure 5-.**
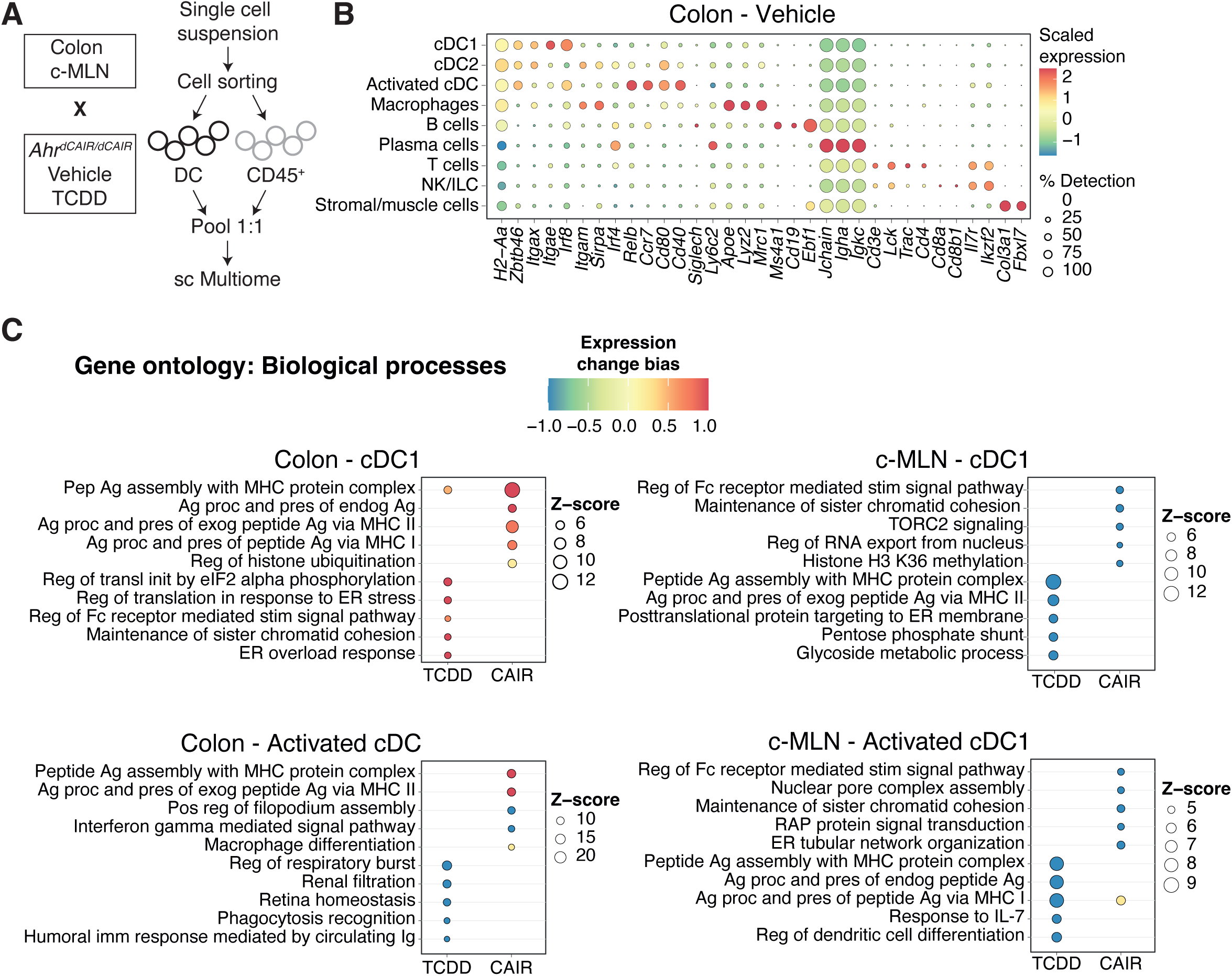
Single cell RNAseq analysis. (**A**) Experimental setup for sc Multiome from dendritic cells and CD45+ cells. **(B)** Dot plot showing the putative marker genes across cell types defined in the vehicle sample in the Colon. The size of the dot represents the percentage of cells within clusters expressing the gene, while the colour scale represents the average expression level in that cluster. **(C)** Gene ontology enrichment analysis of biological processes for differentially expressed genes in TCDD and *Ahr^dCAIR/dCAIR^* mice compared to vehicle for the indicated cell type. The colour scale represents the expression change bias, which represents the proportion of upregulated and downregulated genes that belong to that pathway, while the size of the dot shows the Z-score. Differentially expressed genes were defined as those with an adjusted p-value <0.01 and enriched pathways have a p-value <0.01.

Clusters of the major immune cells and their different subpopulations were visualised using uniform manifold approximation and projection (UMAP) of the RNA-seq datasets. UMAPs generated for the RNA-seq and ATAC-seq datasets from colon and c-MLN of all conditions showed a similar representation of cell types (Fig. EV5A,B). Fig.5B and Fig.EV5C give an overview of marker expression for identification of the major immune cell types found in colon and c-MLN, respectively.

To identify the pathways modulated by *Ahr^dCAIR/dCAIR^*mice and TCDD in cDC we performed gene ontology analysis using differentially expressed genes (DEG) in each condition compared to the vehicle group. Fig.5C and Appendix Fig.S3 give an overview comparing TCDD to *Ahr^dCAIR/dCAIR^* mice in dendritic cells of colon and c-MLN. The most prominent difference was seen in pathways corresponding to antigen processing and presentation which was markedly enhanced in dendritic cell subsets from *Ahr^dCAIR/dCAIR^* mice particularly in the colon compared with vehicle control and dendritic cells from TCDD treated mice, where genes in these pathways were down-regulated, particularly in c-MLN. The populations of activated cDC1 and cDC2 characterised by high levels of *Ccr7*, *Cd40* and *Relb*, were not well separated in the colon and were therefore considered together.

There was no evidence of increased antigen processing and presentation pathways in c-MLN for *Ahr^dCAIR/dCAIR^*except in the subset of activated cDC1. Thus, the transcriptomic profiling confirmed the deficiency in antigen processing and presentation by dendritic cells from TCDD treated mice, but highlights enhancement of this function in *Ahr^dCAIR/dCAIR^* mice above even vehicle treated mice. This clearly indicates that prolonged AHR activation itself is not directly linked with suppression of the immune response.

### Chromatin accessibility differences between TCDD treated and *Ahr^dCAIR/dCAIR^* mice

Ontology of differentially expressed genes and neighboring regions with differentially accessible regions identified by ATAC-seq in dendritic cells from TCDD treated compared to *Ahr^dCAIR/dCAIR^* mice indicated small but discernible differences in genes linked to dendritic cell function. Examples for this are shown in Fig.6 where shaded areas highlight the peaks in question in colon cDC1 and activated cDC1 in the c-MLN, while violin plots show gene expression. Reduced accessibility and gene expression was seen in DC from TCDD treated mice for amino acid transporter *Slc7a5* that plays and important role in the TORC1 pathway and influences DC/T cell crosstalk, affecting maturation, migration and antigen processing (Shao *et al*, 2024) as well as for *Hspa1b,* which encodes a heat shock protein that functions as chaperone with a role in the cellular stress response and protein quality control, including a role in antigen processing and presentation. Similarly *H2-DMb1*, a gene located in the Major Histocompatiblity class II locus that affects peptide binding to MHC class II in conjunction with chaperone co-factors was less accessible in cDC1 from TCDD treated mice. The final example is *Actb* encoding actin, vital for synapse formation between T cells and dendritic cells as well as functions such as migration and antigen capture.

**Figure 6-.**
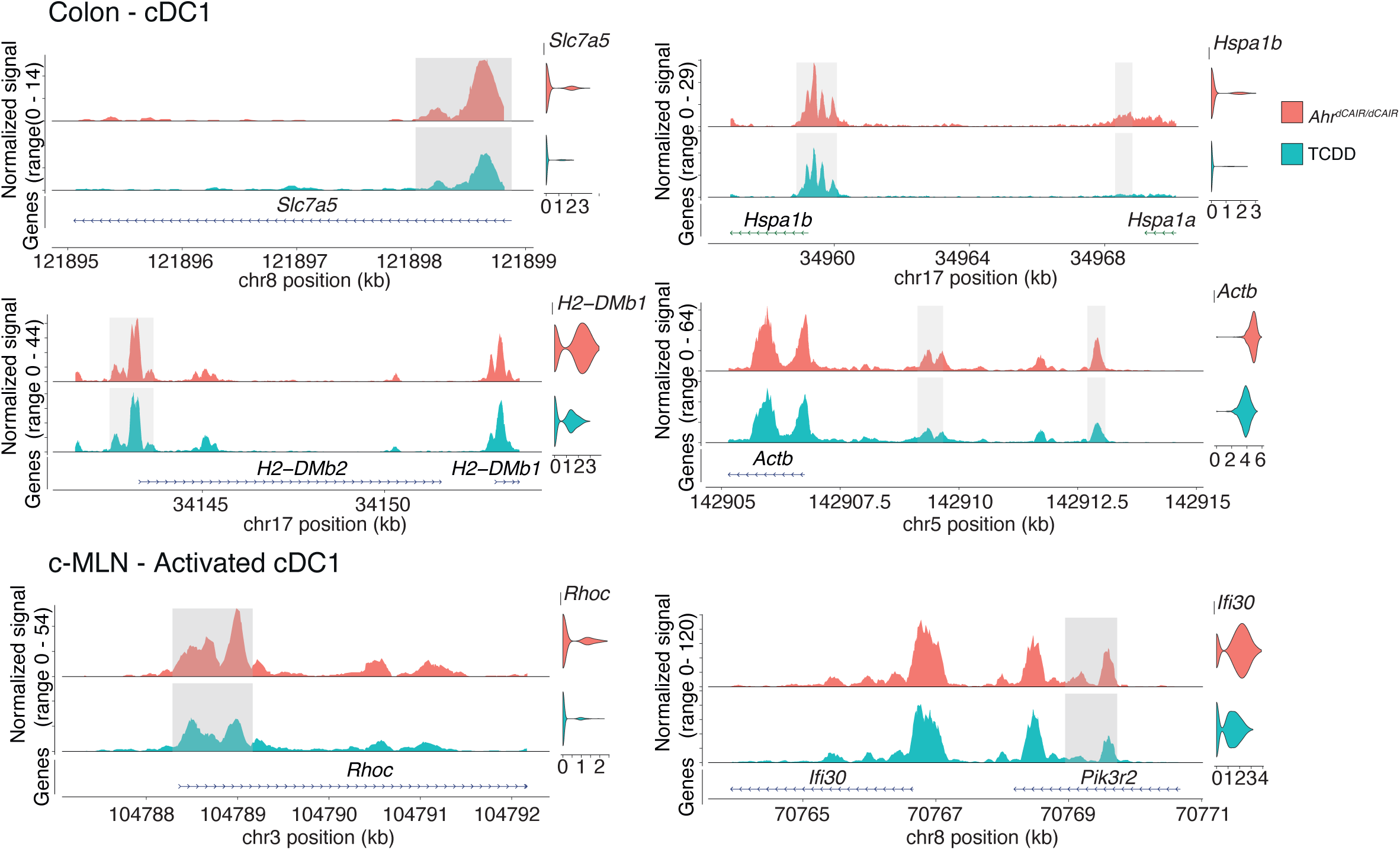
Gene expression and chromatin accessibility in cDC1 of *Ahr^dCAIR/dCAIR^* mice compared to TCDD. ATACseq signal profiles of differentially accessible peaks and linked genes that are also differentially expressed in cDC1 populations in colon and MLN. Differentially accessible peaks are shaded in grey, while violin plots show gene expression in the respective population and condition. ATAC-seq profiles were generated from pseudo-bulk chromatin accessibility data and violin plots show normalized expression values at the single cell level.

In activated cDC1 in c-MLN *Rhoc*, a member of the Rac subfamily of Rho GTPases which are important for migration and adhesion as well as the reorganisation of the actin cytoskeleton (Ridley, 2006) was differentially accessible as was *Ifi30,* a lysosomal thiol reductase that plays a role in MHC class II restricted antigen processing.

Overall the ATAC-seq data confirm that TCDD interfered with the induction of a group of genes that are required for crucial functions of dendritic cells such as migration, maturation, activation and antigen presentation.

## Discussion

The increasing evidence for involvement of environmental factors in many inflammatory diseases has led to an interest in AHR as a potential therapeutic target. However, there is still a lack of clarity on what defines AHR agonists with beneficial properties compared with those that could cause toxicity and other adverse effects, which hampers progress with AHR focused interventions.

The prevailing rationale for AHR-mediated toxicity seen at exposure to TCDD and other persistent xenobiotic AHR ligands has been associated with their prolonged AHR activation compared to rapidly metabolised endogenous ligands. The argument is persuasive as closer scrutiny of the AHR pathway highlights the emphasis on induction of negative feedback regulation following activation, such as the induction of Cyp1 family enzymes as well as the AHR repressor (AHRR). Thus, one could assume that tight transient activation of the pathway is paramount for its physiological functioning. In this study we tested whether prolonged activation of AHR per se is an inducement of detrimental activity that would perturb the normally beneficial AHR function in an intestinal infection. For this we used TCDD as well as a knockin model (*Ahr^dCAIR/dCAIR^*) for constitutively active AHR under full control of the endogenous AHR locus (Ye *et al*., 2017). *Ahr^dCAIR/dCAIR^* mice showed constitutive expression of *Cyp1a1* indicative of AHR activation, but no overt adverse effects were noticeable under steady state conditions. This is in contrast to a previously published model of constitutive AHR activation in which a construct lacking the PAS-B domain was expressed as a transgene under a heterologous promoter, resulting in the development stomach cancer (Andersson *et al*., 2002). The *Ahr^dCAIR/dCAIR^* mice did however replicate the liver- and thymus-toxicities that have been reported following exposure to TCDD.

Given that the intestinal microenvironment is substantially influenced by AHR activity in multiple cell types, we chose the intestinal infection model with *C. rodentium* to probe the consequences of either TCDD application or endogenous constitutive AHR activity.

Against our initial assumption, *Ahr^dCAIR/dCAIR^* mice showed no defects during this infection and dealt with it more efficiently than wildtype mice, exhibiting lower bacterial levels, less inflammation and increased production of IL-22, a major determinant of barrier protection in this infection (Basu *et al*., 2012; Li *et al*, 2018; Melchior *et al*, 2024). In contrast, mice exposed to TCDD prior to infection clearly had a suboptimal response with delayed clearance and compromised antibody responses. Notably, both TCDD treated mice as well as *Ahr^dCAIR/dCAIR^* mice were more resistant to DSS induced colitis, showing markedly reduced pathology. As the DSS model primarily affects the epithelium, this could be attributed to preferential activity in colonic epithelial cells where AHR activity is normally markedly lower than that seen in the small intestine (see Fig.2C and (Zhou *et al*, 2023)). It therefore seems that enhancement of AHR activity in the colon might be beneficial for the organism even if it involves exposure to xenobiotic TCDD.

Nevertheless, our data indicate that TCDD impairs the response of immune cells participating in the defense against *C. rodentium*. This has been observed in other organs, notably the lung, and the toxicology literature refers to TCDD as an immunotoxic substance (Kerkvliet, 2002). Although the impairment of the B cell response to *C. rodentium* by TCDD was not extensive, it did cause a delayed clearance of bacteria. The role of B cells in the host defense against *C. rodentium* is recognised but that of specific Ig isotypes is somewhat controversial (Belzer *et al*, 2011; Maaser *et al*., 2004). Here, the two Ig subclasses that were reduced following exposure to TCDD were IgG2b and IgG3, which are important for complement fixation and opsonisation of bacteria. IgG bound to *C. rodentium* virulence factors leads to their selective elimination by neutrophils (Kamada *et al*, 2015) and so it is conceivable that a reduction in this type of antibody may compromise speedy clearance of the pathogen. Furthermore, these two Ig subtypes are produced mainly by B1 cells and marginal zone B cells and are known to react against gram negative bacteria (Cerutti *et al*, 2013; Zeng *et al*, 2016). These cells also express the highest levels of AHR (Fig.1 and (Villa *et al*, 2016)). While it has been suggested that repression of the differentiation factor Blimp-1 by TCDD-activated AHR underlies the suppressive effect of TCDD on B cell responses (De Abrew *et al*., 2011), this explanation does not fit with data showing that IgG3 producing B1 B cells are largely Blimp-1 independent (Savage *et al*, 2017). It was notable that the numbers of dendritic cells and B cells were markedly decreased in TCDD treated mice 14 days after infection, probably reflecting more a failure to expand rather than increased cell death. This is supported also by the observation that the lymph nodes of TCDD treated mice were considerably smaller than those of vehicle treated mice.

In order to get more mechanistic insight into the immune cell affected by TCDD and the comparison to constitutive AHR activation, we decided to carry out single cell RNA sequencing combined with ATAC sequencing of cells isolated from colon and c-MLN on day 5 after infection. This is a timepoint when the adaptive immune response to infection is beginning to take over from the initial innate response (Ahlfors *et al*, 2014; Zindl *et al*., 2022). To our knowledge, this is also the first single cell RNAseq characterization of the immune compartment following *C. rodentium* infection, following a previous effort focused solely on CD4+ T cells (Kiner *et al*, 2021).

We enriched for dendritic cells to facilitate their profiling as they are normally present in very low numbers. Their obligatory role for T cell activation supporting humoral as well as cellular immunity is well known, but it is becoming increasingly clear that B cells also may directly communicate with dendritic cells for effective antibody responses (Steiner *et al*, 2022). Our data highlight the markedly increased dendritic cell function in antigen processing and presentation seen in *Ahr^dCAIR/dCAIR^* mice in comparison with TCDD or vehicle treated mice. This was striking in cells isolated from colon but not the c-MLN with the exception of a population of activated dendritic cells, potentially indicating that most dendritic cells that had taken up and processed antigens in the colon had not yet migrated to the colon draining lymph nodes.

The overlay of ATAC-seq and RNA-seq also confirmed that chromatin loci close to genes involved in migration, activation and antigen processing/presentation were more accessible in dendritic cells from *Ahr^dCAIR/dCAIR^* than in those of TCDD treated mice, coinciding with higher gene expression.

Taken together our data confirm previous data in the literature showing that TCDD results in suppression of immune responses. However, they also clearly indicate that prolonged AHR activation per se cannot account for this, as *Ahr^dCAIR/dCAIR^* mice with constitutive AHR activity showed such profound enhancement of precisely those parameters that were suppressed in TCDD treated mice. It remains to be shown mechanistically what the underlying cause for the immunosuppressive effect of TCDD might be. Prolonged activation of AHR appears linked to known parameters of toxicity evident in liver and thymus, but in contrast may even enhance immune reactivity. It should be interesting to investigate how *Ahr^dCAIR/dCAIR^* mice respond to different kinds of immune challenge.

### Reagents and Tools table

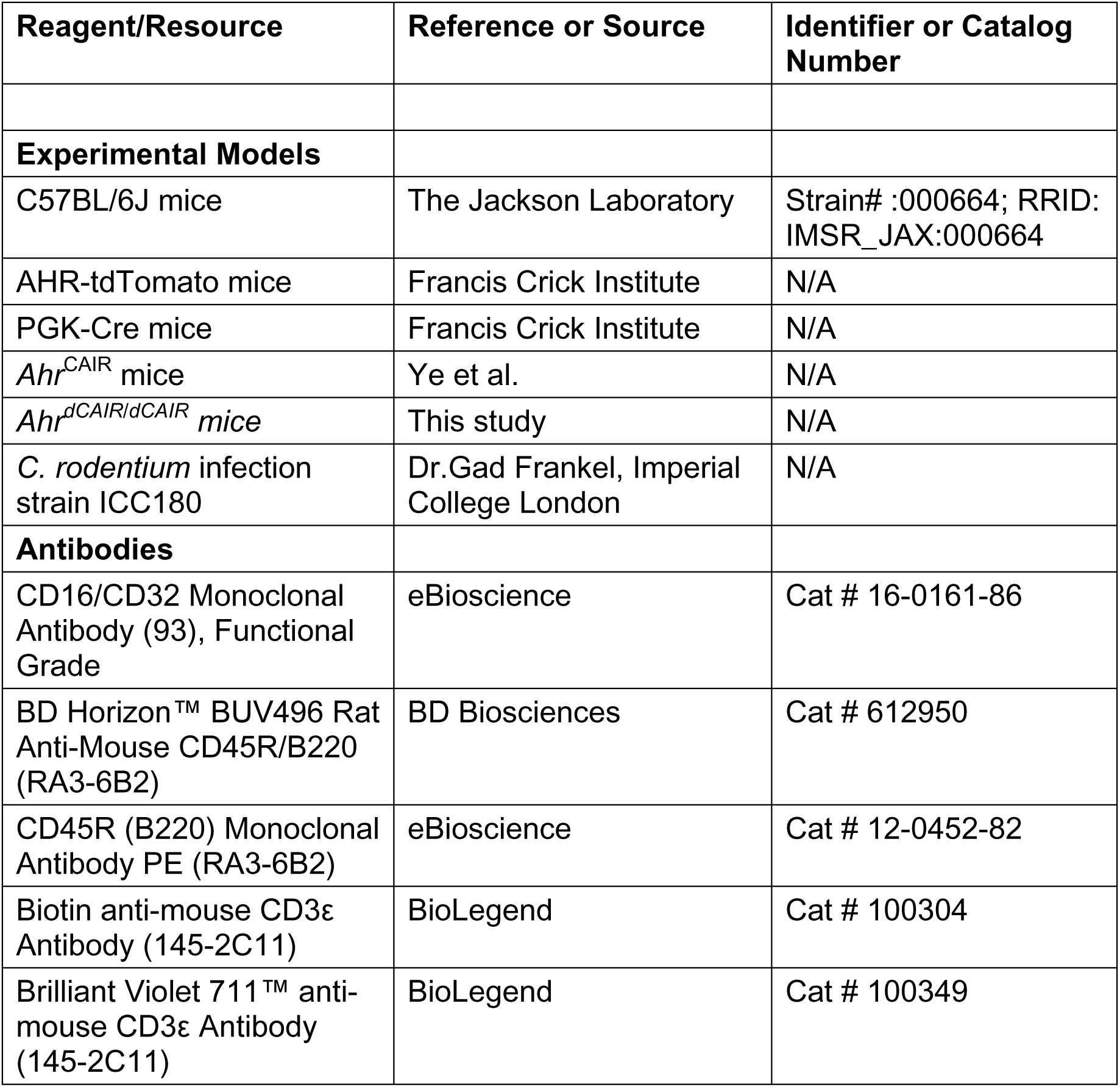
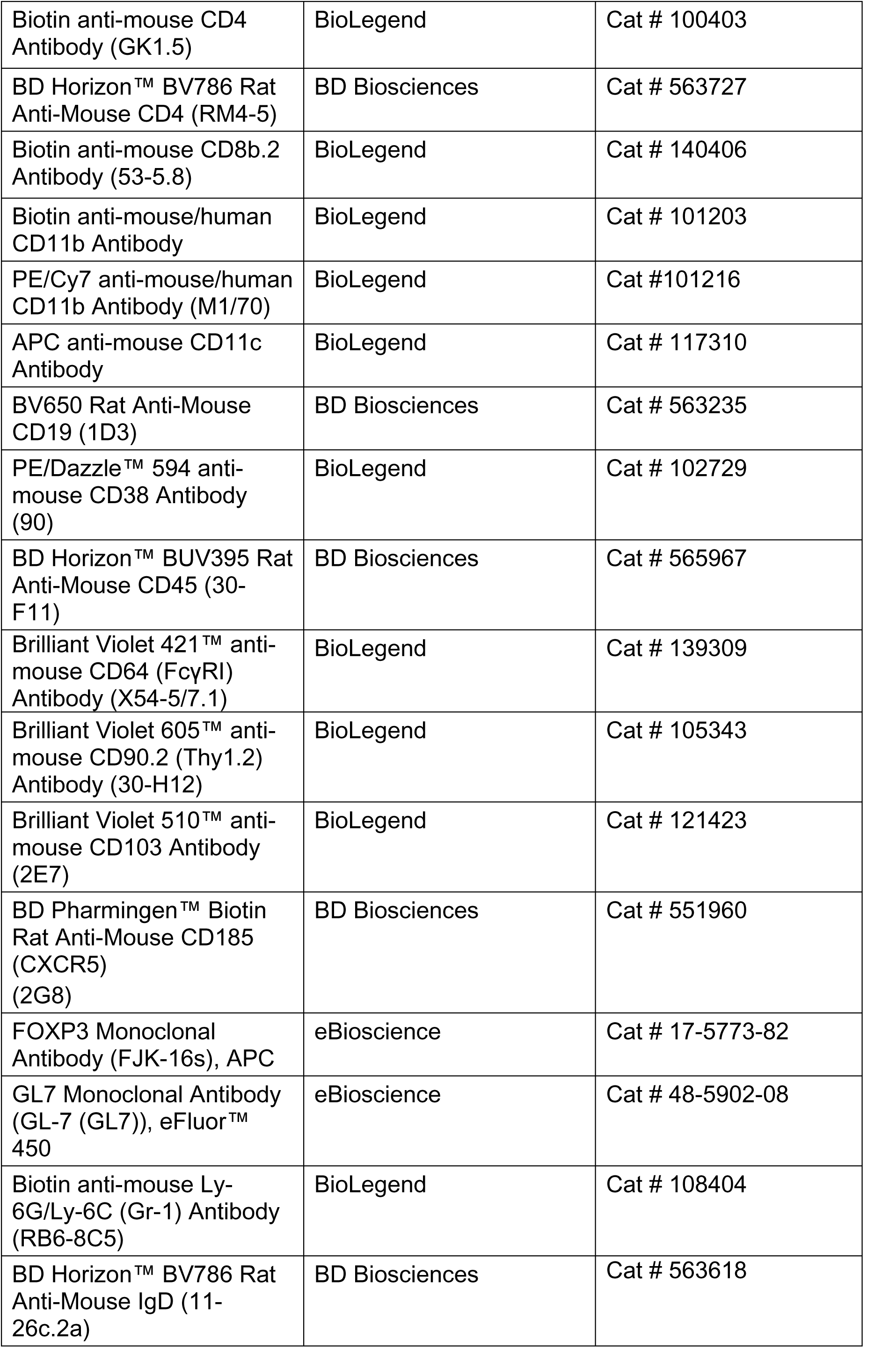
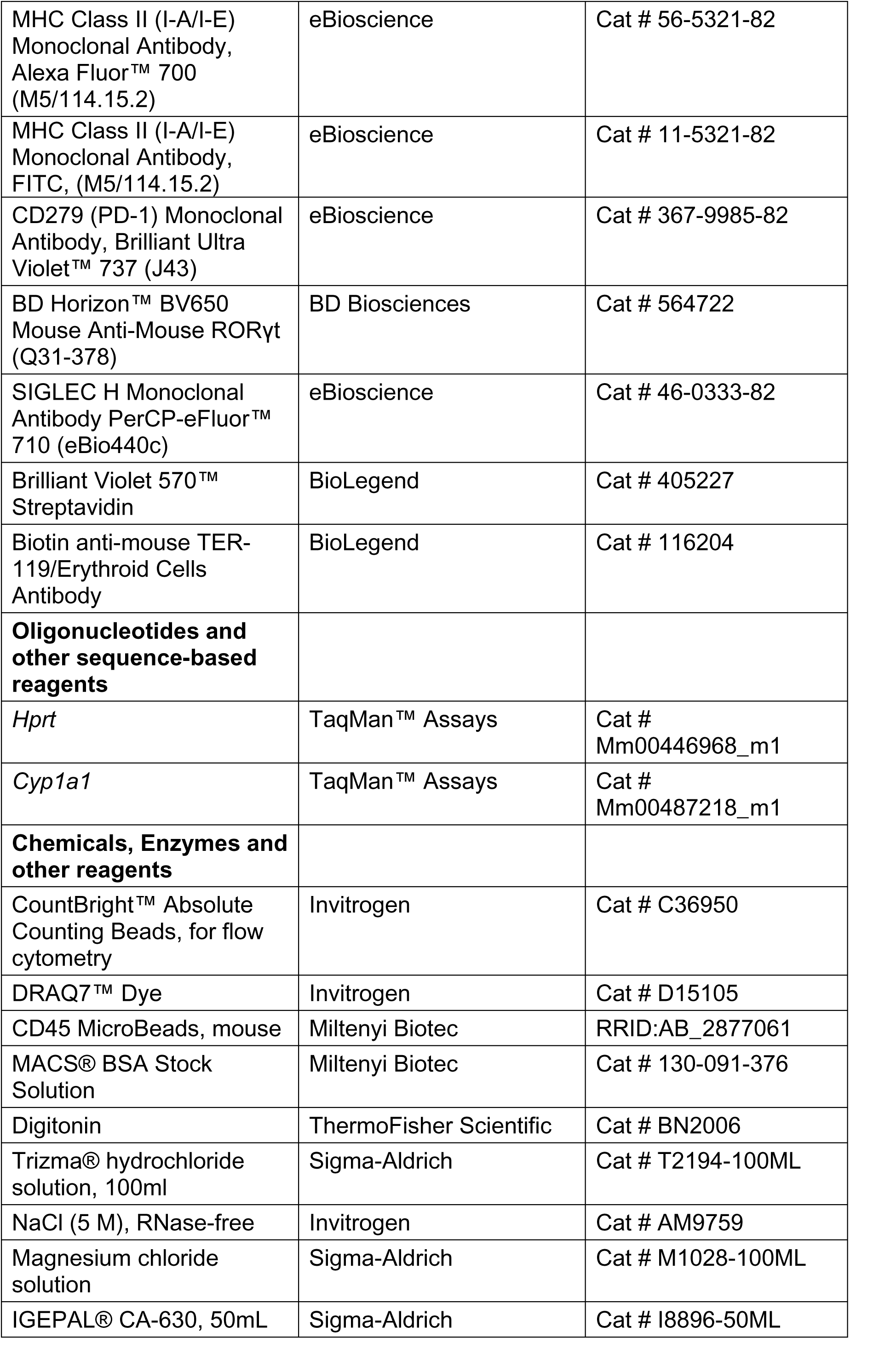
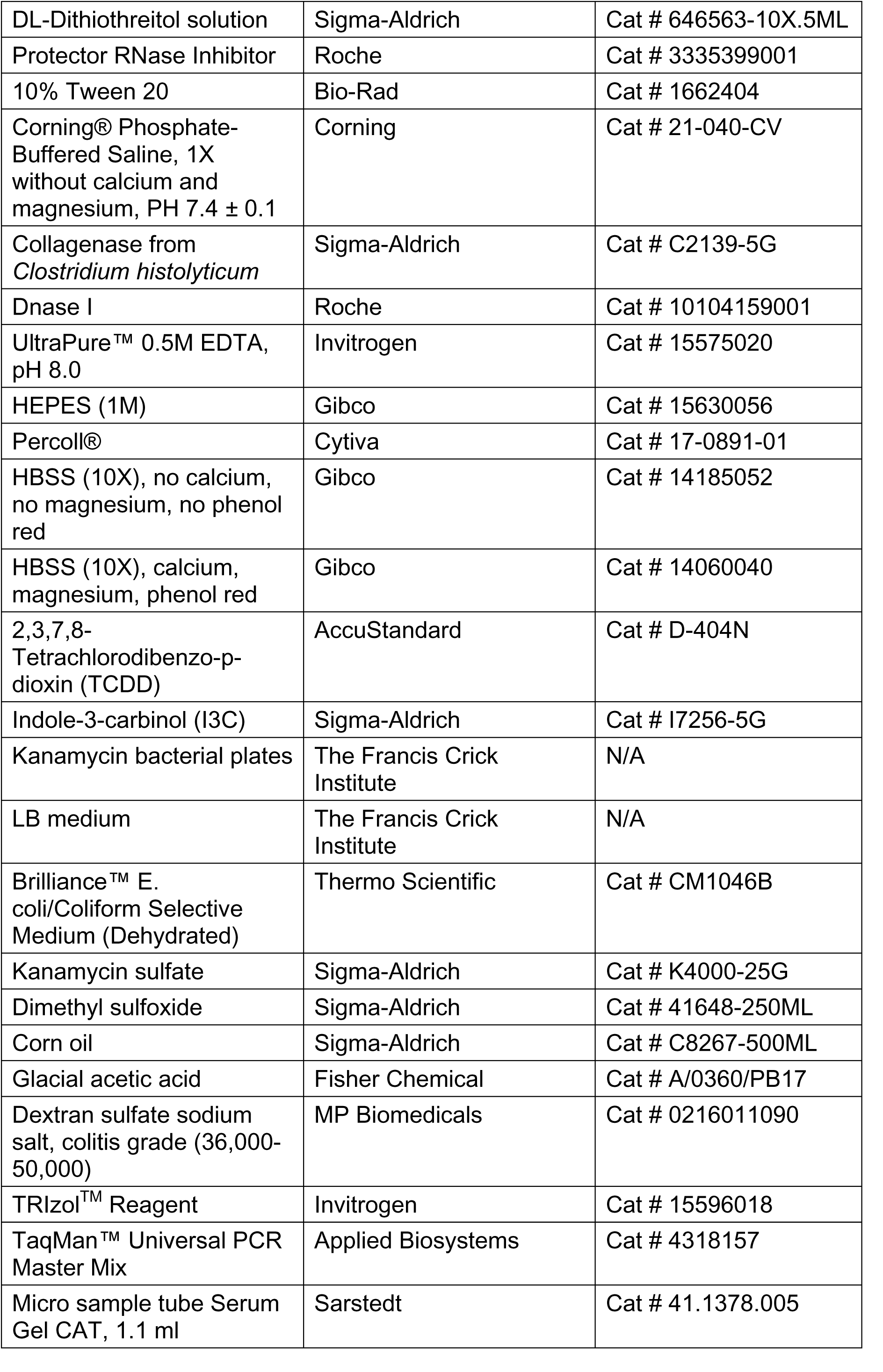
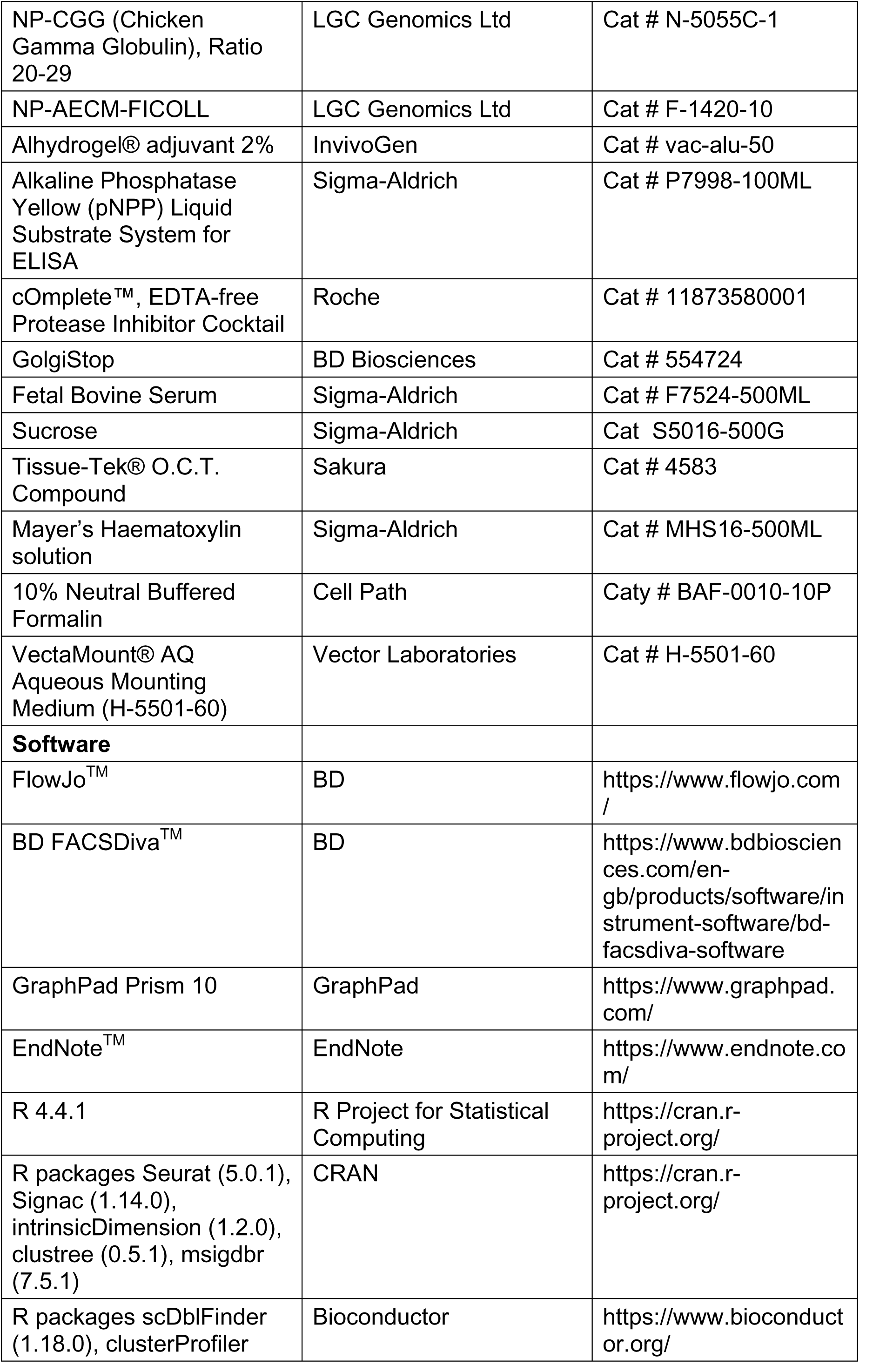
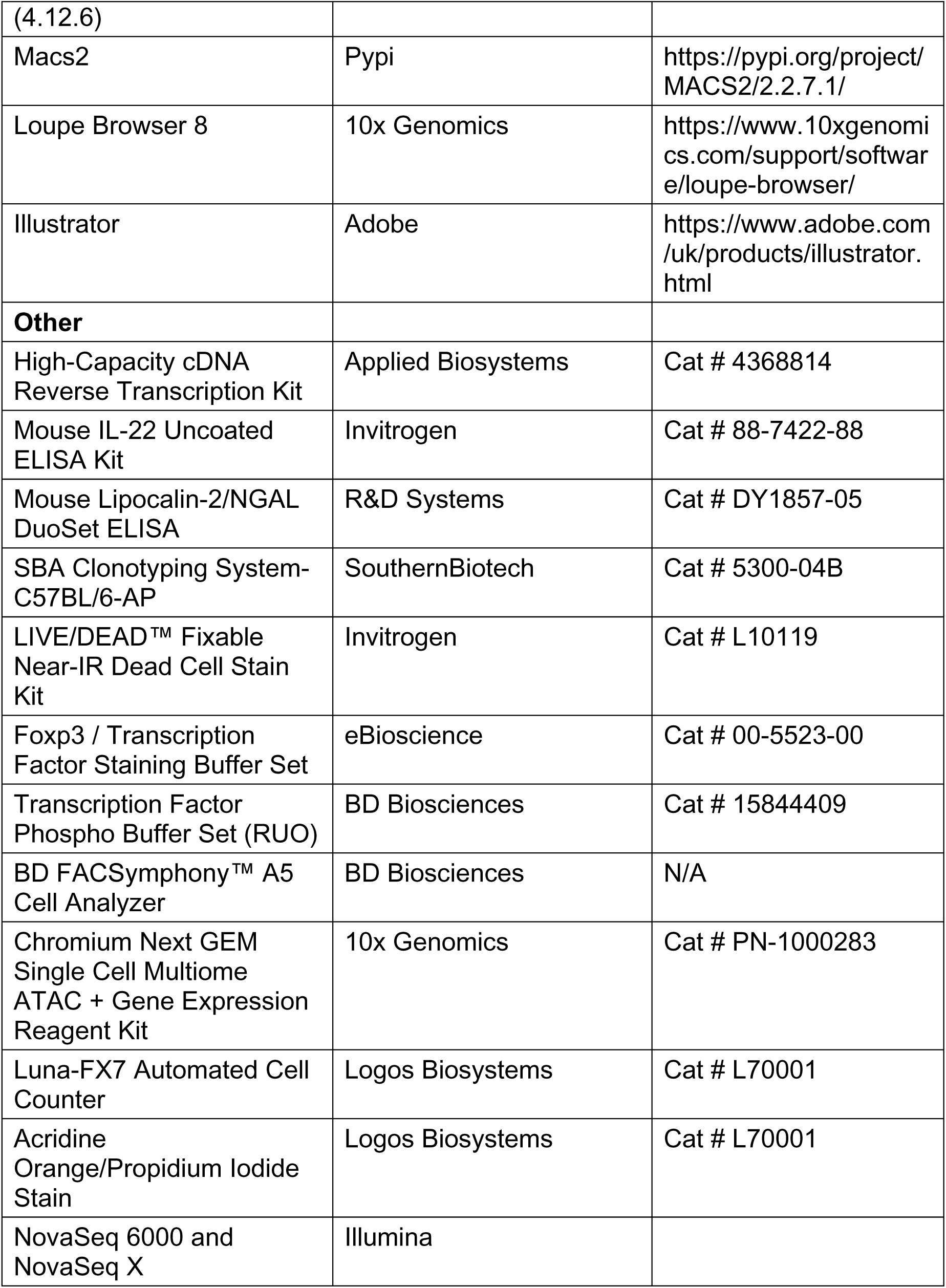

## Methods and Protocols

### Mice

AHR-tdTomato, PGK-Cre, *Ahr^CAIR^*, *Ahr^dCAIR/dCAIR^* and C57BL/6J mice used in this study were bred and maintained in individually ventilated cages under specific pathogen-free conditions at the Francis Crick Institute, according to protocols approved by the UK Home Office and the Ethics committee of the Francis Crick Institute. Mouse experiments were conducted according to the guidelines detailed in the Project Licenses granted by the UK Home Office to Brigitta Stockinger (PP0858308). Mice were age- and sex-matched and between 6-9 weeks old when first used. Both female and male mice were used. Exclusion criteria such as inadequate staining or low cell yield owing to technical problems were pre-determined. Mice were randomly assigned to experimental groups.

### Infection with Citrobacter rodentium

For C. rodentium infection, a frozen stock of C. rodentium strain ICC180 (kindly provided by Dr. Gad Frankel, Imperial College London) was streaked overnight on a Luria-Bertani (LB) agar plate supplemented with 50 μg/ml Kanamycin and incubated overnight in a dry incubator at 37 °C. A single colony was picked and grow in LB broth supplemented with 50 μg/ml Kanamycin and grown to log phase, followed by centrifugation and resuspension in PBS. Mice were orally gavaged with 200 μl of PBS containing 2 x 10^9^ *C. rodentium* CFU. To determine the bacterial load, tissue pieces or faecal pellets were weighed and homogenized in sterile PBS and serial dilutions were plated onto LB plates supplemented with 50 μg/ml Kanamycin or plates with Brilliance *E. coli*/Coliform Selective Medium and incubated overnight at 37 °C. The numbers of CFU were normalized to the weight of the fecal pellets.

Sample sizes for experimental groups were determined based on available litter sizes or previous experience in the lab with the *C. rodentium* model. Animals that on day 7 post infection had pathogen levels below 10^7^ CFU per gram of tissue were excluded from the experimental results.

### Ligand administration

TCDD (AccuStandard) was dissolved at 100 μg/ml in DMSO by sonication for 30 minutes and stored at −80 °C. For *in vivo* administration, TCDD was dissolved in corn oil at a concentration of 2 μg/ml and administered by oral gavage as a single dose of 10 μg/kg of body weight. To test the effect of TCDD on the clearance of *C. rodentium*, a single dose of 10 μg TCDD per kg was given 6 days before the infection. Vehicle treated mice were given an equivalent dose of corn oil with 2% DMSO at a volume of 5 μl.

For *in vivo* administration, I3C (Sigma-Aldrich) was dissolved at a concentration of 50 mg/ml in 10% DMSO, 0.5% glacial acetic acid, and 89.5% corn oil. I3C was administered by oral gavage as a single dose of 250 mg/kg of body weight.

### Chemical tissue extraction

Extraction of TCDD was perfomed using a sequence of extraction and precipitation steps depending on type of tissue. Pre-weighed intestinal tissues were homogenized frozen using pre-chilled 5mm stainless steel beads and pre-chilled tissue-lyser II (Qiagen, Germany) at 30Hz, in 30s intervals. Liver and adipose tissues were homogenized in a similar way, but not frozen. Isopropanol was added to the homogenates of ground tissues, and re-homogenizedfor 30s. Internal standard (PCB118) was added and the homogenates extracted by vortexing, followed by centrifugation at 5000 rpm for 1 min. The isopropanol phase was transferred to a clean tube, and the pellet re-extracted with 3-methylpentane (3MTP) by vortexing, followed by centrifugation at 5000 rpm, and tranfer of the 3MTP supernatant to the isopropanol phase. Water containing 0.1M sulphuric acid was added to the combined supernatants and the suspension was centrifuged. The organic phase was transferred to a clean tube and concentrated sulphuric acid was added, followed by gentle mixing and centrifugation. The upper phase was finaly transferred to an HPLC vial and stored at 4 °C until analysis. Due to their low concentrations of TCDD, the replicate samples of intestinal tissue extracts were pooled and run a through sulphuric acid/silica gel column as an additional clean-up step before being analysed as described below. Serum samples were extracted the same way, starting from the step adding isopropanol. Extraction of I3C and ICZ is described in the Appendix Supplementary Methods.

### Chemical analysis

Full description of the chemical analysis can be found in the Appendix Supplementary Methods. Briefly, tissue levels of TCDD were determined by GC-MS analysis, with the MS operated in electron capture negative ionization (ECNI) mode, and separation was achieved on a J&W DB-5MS UI capillary column (30 m × 0.25 mm i.d. × 0.25 µm film thickness; J&W Scientific, CA, USA) and detection was done in selective ion monitoring mode (SIM). The internal standard PCB118 was detected using the ion m/z: 327.9 while TCDD was detected using the chlorine ions m/z: 35 and 37. Quantifications were conducted against external calibration curves.

Analysis of I3C and ICZ were conducted using a Bio compatible HPLC system and separation was achieved using a Poroshell 120 EC-C18 column (3.0 x 50 mm, 2.7-micron particle size, Agilent InfinityLab, Agilent, US). I3C, ICZ, and the internal standard FICZ were mionitored by both UV detection (280/254 nm) and fluorescence detection (390:518 nm excitation/emission), and quantified against external calibration curves.

### Liver and thymus toxicity analysis

6-7 week old mice were given corn oil or a single dose of 10 μg/kg of body weight and analyzed 6 days later. Mice were weighed before organ collection. Thymi were cleaned and weighed. Whole livers were collected and weighed and a half of the left lobe was fixed in 10% neutral-buffered formalin overnight. After washing twice in PBS, the organs were cryoprotected in 30% sucrose overnight. Tissues were embedded in OCT and stored at −80C. For the Oil Red O staining, 10 μm sections were fixed in 10% neutral buffered formalin, rinsed twice in isopropanol and stained in freshly prepared Oil Red O working solution (3g/60ml isopropanol:40ml distilled water). Slides were rinsed in isopropanol, counterstained with Mayer’s Haematoxilin (Sigma-Aldrich) and mounted using aqueous mounting medium H5501 (Vector Laboratories).Randomized images were obtained using a Zeiss Axioscan Z1 slide scanner. From each liver section, 5 smaller images were used for quantitative analysis using Fiji, as described in (Graelmann *et al*, 2024). Briefly, the intensity of the red signal was calculated and the optical density was estimated normalizing the mean intensity on the maximal intensity.

### DSS colitis

For induction of DSS colitis, mice were provided with 2% w/v Dextran sulfate sodium (DSS) (MP Biomedicals) in their drinking water *ad libitum* for 5 days, followed by normal drinking water for 25 days.

### Estimation of Lipocalin-2 levels

Fecal lipocalin-2 levels were measured in stool homogenates using the mouse Lipocalin-2/NGAL DuoSet ELISA (R&D Systems) according to the manufacturer’s instructions and values were normalized to the weight of the fecal pellets.

### Cell isolation

Colon and small intestine were cleaned of luminal contents, cut open longitudinally and washed in PBS. The epithelial layer was removed by incubation in HBSS without Ca^2+^ / Mg^2+^ (Gibco), 5% FBS and 2 mM EDTA (Invitrogen) for 40 min at 37 °C at 200 rpm. Intestinal pieces and the c-MLN were washed in PBS, cut into small pieces and digested with 1.5 mg/ml Collagenase VIII (Sigma-Aldrich), 50 μg/ml Dnase I (Roche) with 5% FBS (Sigma-Aldrich), 15 mM HEPES (Gibco) in Ca^2+^ / Mg^2+^-free HBSS for 30 minutes. Cells were filtered through a 100-μm cell strainer, washed in PBS with 5% FBS, filtered through a 70-μm strainer and washed again.

### Flow cytometry

Cell suspensions were prepared as described for the indicated organ and incubated with anti-CD16/32 (eBioscience) and LIVE/DEAD™ Fixable Near-IR Dead Cell Stain Kit (Invitrogen) for 15 min at 4°C and washed in PBS. For surface stainings, cells were incubated with directly conjugated antibodies in PBS with 5% FBS at 4 °C for 20 minutes. Cells were washed in PBS with 5% FBS and optionally fixed in 4% formaldehyde in PBS for 45 min to 18h at 4 °C. For intracellular staining of transcription factors or cytokines, cells were fixed and permeabilized with the Foxp3 / Transcription Factor Staining Buffer Set (eBioscience), according to the manufacturer’s instructions. For intracellular stainings of cells obtained from AHR-tdTomato mice, cells were fixed and permeabilized using the Transcription Factor Phospho Buffer Set (RUO) (BD Biosciences), according the the supplied protocol. Samples were acquired following standard procedures on a BD FACSymphony™ A5 Cell Analyzer (BD Biosciences). Acquired data was analyzed using FlowJo software (https://www.flowjo.com/). B cells were gated as Live CD45+ Lin- (CD3, CD4, CD8, F4/80, CD11b, Gr1, Ter119) and analyzed as follows: B1 (CD19+ B220-), B2 (CD19+ B220+), GL7+ CD38-B cells (GL7+ CD38-CD19+ B220+), IgD-GL7+ B cells (IgD-GL7+ CD19+ B220+). Myeloid cells were gated as Live CD45+ and defined as: dendritic cells (CD64-CD11c+ MHCII+), CD11b+ DCs (CD11b+ CD64-CD11c+ MHCII+), CD103+ DCs (CD103+ CD64-CD11c+ MHCII+), CD11b+ CD103+ DCs (CD11b+ CD103+ CD64-CD11c+ MHCII+), macrophages (CD64+ MHCII+), pDCs (CD11clo MHCII-SiglecH+ B220+). ILCs and T cells were gated as Live CD45+ CD90+ and analyzed as follows: ILC (CD3-), T cells (CD3+), CD4+ T cells (CD4+), Foxp3+ CD4+ T cells (CD4+ Foxp3+), Rorgt+ CD4+ T cells (Rorgt+ CD4+), Tfh cells (PD-1+ CXCR5+), CD8+ T cells (CD8a+).

Experimental groups were masked during the preparation of the cell suspension and data analysis.

### Colon explant cultures

Colon tissue pieces (0.5-1cm length) were weighed and cultured for 24h at 37 °C in complete IMDM medium. IL-22 cytokine levels in the supernatants were determined by ELISA (Invitrogen) and concentrations were normalized to the weight of the explants.

### RNA isolation, cDNA synthesis and and qRT-PCR

RNA was isolated from intestinal and white adipose tissue using TRIzol™ Reagent (Invtitrogen), according to the manufacturer’s protocol. cDNA was synthesized with the High-Capacity cDNA Reverse Transcription Kit (Applied Biosystems) and real-time quantitative PCR was performed using the TaqMan™ Universal PCR Master Mix (Applied Biosystems) and respective probes. mRNA expression were determined using the ΔCt method, relatively to hypoxanthine-guanine phosphoribosyltransferase (*Hprt*) gene expression.

### NP-Ficoll and NP-CGG immunizations and measurement of anti-NP antibodies

NP-Ficoll or NP-CGG (LGC Genomics) was dissolved in PBS and mixed with Alhydrogel (InvivoGen) at a 1:1 volume ratio. Mice were immunized with 10 μg NP-Ficoll or NP-CGG administered as a volume of 100 μl intraperitoneally. Fourteen days after immunization, whole blood was collected in Micro sample Serum Gel tubes (Sarstedt) and serum was prepared by centrifugation at 10000xg for 5 minutes.For measuring anti-NP antibodies, pleates were previously coated with 10 μg/ml NP-Ficoll or NP-CGG overnight at 4 °C (Akkaya *et al*, 2017). Coated plates were washed in PBS plus 0.05% Tween 20, blocked in PBS plus 1% BSA and washed before the addition of a 5-fold serial dilution of sera. Samples were incubated for 2h at room temperature. Ig isotypes were detected with goat anti-mouse IgG1, IgG2b, IgG2c, IgG3 antibodies conjugated to alkaline phosphatase (SouthernBiotech) diluted in PBS + 1% BSA and incubated for 1 hour at room temperature. After washing, plates were developed with *p*-nitrophenylphosphate (pNPP) substrate and read in a Tecan plate reader at OD_405_.

### Anti-*C. rodentium* Ig ELISA

Analyses were performed on fecal lysates or serum prepared from whole blood collected in Micro sample Serum Gel tubes (Sarstedt) and by centrifugation at 10000xg for 5 minutes. Heat-killed *C. rodentium* was prepared as previously described (Bry & Brenner, 2004). Briefly, an 18-hour culture was resuspended in PBS plus a cOMPLETE Protease Inhibitor cocktail (Sigma-Aldrich) up to a volume with an OD_600_ of 1.0. The culture was heat-killed by incubation at 60 °C for 1h, aliquoted and frozen at −80 °C until further use.

For the ELISA aliquots were diluted 50 times in PBS for coating the plates, and stored overnight at 4 °C. Anti-*C.rodentium* antibodies were detected using goat anti-mouse IgA, IgG1, IgG2b, IgG2c and IgG3 antibodies conjugated to alkaline phosphatase (Southern Biotech), following the protocol described in the above section.

### Single-cell multiome sequencing of immune cells from the c-MLN and colon

Four to six mice per group received an oral administration of with vehicle TCDD 6 days prior to infection with 2 x 10^9^ CFU *C. rodentium* as as described above. At day 5 post infection, cells from the c-MLN and colon lamina propria were isolated as described above and pooled in one sample of the respective organ and group. The immune fractions were enriched using CD45 MicroBeads according to the manufacturer’s protocol. Cells were stained with CD45, CD11c, MHCII and CD64 surface antibodies and DRAQ7 Dye to exclude non-viable cells, and then sorted on a BD FACS Aria Fusion into a 1.5ml tube with PBS + 5% FCS using a 100 μm nozzle and a sorting precision protocol 16-16-0. Immune cells (Live CD45+) and dendritic cells (Live CD45+ CD11c+ MHCII+ CD64-) were sorted individually and mixed at a 1:1 ratio, for a total of 60000-160000 cells per sample. Cell nuclei were isolated according to the Demonstrated Protocol: Nuclei Isolation for Single Cell Multiome ATAC + Gene Expression Sequencing (CG00053 – Rev C, 10x Genomics) with the following modifications: nuclei for all samples were isolated using the Low Cell Input Nuclei Isolation protocol were cells were centrifuged at 300xg for 5 min at 4 °C and resuspended in 50 μl PBS + 0.04% BSA, lysed in chilled Lysis Buffer for 4 min on ice. After the incubation, wash buffer was added without mixing and the nuclei were centrifuged for 5 min at 500xg at 4 °C. Nuclei were washed once in Diluted Nuclei Buffer and resuspended in Diluted Nuclei Buffer. The concentration and viability of the single nuclei suspension was measured using (1) acridine orange (AO) and propidium iodide (PI) and the (2) Luna-FX7 Automated Cell Counter. Approximately (3) [5,000-10,000] nuclei were transposed, then loaded on Chromium Chip and partitioned in nanolitre scale droplets using the Chromium Controller and Chromium (4) Next GEM Single Cell Reagents (Chromium Next GEM Single Cell Multiome ATAC + Gene Expression Reagent Kits User Guide, User Guide, CG000338). A pool of 736,000 10x Barcodes was sampled to separately and uniquely index the transposed DNA and cDNA of each individual nucleus. ATAC and gene expression (GEX) libraries were generated from the same pool of pre-amplified transposed DNA/cDNA and sequenced using the Illumina (5) NovaSeq 6000 (ATAC) and NovaSeq X (GEX). Sequencing read configuration: 28-10-10-90 (GEX), 50-8-24-49 (ATAC). The 10x Barcodes in each library type are used to associate individual reads back to the individual partitions, and thereby, to each individual nucleus. We aimed for 50k reads per cell for GEX libraries, and 25k reads for ATAC libraries.

### Statistics and data analysis

All statistical analysis were performed with GraphPad Prism software (https://www.graphpad.com). Description of the statistical tests used and the number of replicates are described in the corresponding figure legends. Statistically significant outliers were identified using the ROUT method (Q = 5%) and removed from the plots. Data points and n values represent biological replicates and are indicated in the respective figure legend.

### Quantification of gene expression and chromatin accessibility

Libraries were sequenced using Illumina NovaSeq 6000 or NovaSeq X platforms in accordance with 10x recommendations. Sequencing replicates for both modalities were provided to Cell Ranger ARC version 2.0.1 for quantification and filtering of GEMs for putative cells. The 10x-provided “refdata-cellranger-arc-mm10-2020-A-2.0.0” reference was used with default parameters otherwise.

### Filtering Cell Ranger ARC output

After initial QC of the Cell Ranger ARC output, the data were read into a Seurat object using Seurat (5.0.1) and Signac (1.14.0) to manage the gene expression and chromatin accessibilty assays, respectively. All analysis used R version 4.4.1.

Putative cells were filtered according to the number of UMI and features reported per cell and the proportion of mitochondrial expression. Thresholds for these metrics were determined by visual inspection of the distributions in each dataset of each metric, with consideration give to the number of cells retained after filtering.

Doublets were predicted in each unfiltered dataset using the scDblFinder package (1.18.0). The MLN Vehicle, AHR knockout and TCDD datasets showed distinct clusters of doublets which were removed. Similar clusters were not identified in the other datasets.

The filtering parameters used for each dataset, shown in **Appendix Table S1**, retained between 1,538 and 12,830 (mean 7,201) cells per dataset.

### Gene expression analyses

The gene expression assay was analysed using a Seurat pipeline: NormalizeData, FindVariableFeatures, ScaleData, RunPCA, RunUMAP. The number of PCs used for each dataset was determined by considering visual inspection of the elbow plot, marker gene loadings in principal components and results of intrinsicDimension (1.2.0).

Clusters were defined using the FindNeighbors and FindClusters functions, with a range of resolutions from 0.2-2. Marker genes were used to classify cell types in each dataset at an appropriate resolution, aided by clustree (0.5.1) analysis.

Differentially expressed genes were identified within cell types between conditions using the FindMarkers function of Seurat, with the Wilcoxon rank-sum test and correction using the Bonferroni method. The results were subsequently filtered to select features with minimum representation of 0.8 in the cell type, minimum absolute log2 fold change greater than 0.5 and adjusted p-value less than 1%. Functional enrichment was tested using the enricher function of clusterProfiler (4.12.6), within the biological processes defined by the msigdbr package (7.5.1), and a p-value cutoff of 1%.

For plotting, biological processes including ribosomal genes were not included and lists of genes with very similar names were removed to show a wider variety of affected processes.

### Chromatin accessibility analyses

Peaks were identified within cell types using macs2 (2.2.7.1). Peaks were linked to TSS using the LinkPeaks function of Signac, which was used with default parameters.

Differential accessibility was assessed in unified peaks within cell types between conditions using the FindMarkers function, as above. For comparison between chromatin and gene expression assays, the minimum representation in cell type thresholds were relaxed to 0.2 and 0.05 for genes and peaks, respectively. Links between differentially expressed and differentially accessible features were identified from the results of LinkPeaks, with a maximum distance of 10 kb between the peak and TSS.

## Data Availability

Single nucleus RNA-seq and ATAC-seq datasets produced in this study are deposited in the Gene Expression Omnibus (GEO) under accession GSE297700. Supplementary AHR KO snMultiome datasets from both colon and c-MLN are made available (data not shown). Raw and processed data, including interactive Loupe Browser files, are provided for all datasets.

## Acknowledgements

This work was supported by the Francis Crick Institute, which receives its core funding from Cancer Research UK, The UK Medical Research Council, and the Wellcome Trust (FC001159). It was further supported by Wellcome Trust Grant 210556/Z/18/Z to B.S. E.W. is supported by the Swedish research council VR (2020-03418). L.Z. is an Investigator in the Pathogenesis of Infectious Disease and is supported by the Burroughs Wellcome Fund. This work was supported by the National Institutes of Health grants AI132391 and AI157109 (to L.Z.).

For the purpose of Open Access, the author has applied a CC BY public copyright licence to any Author Accepted Manuscript version arising from this submission.’

We would like to acknowledge the Science technology platforms at the Francis Crick Institute. We thank the Biological Research Facility for breeding and maintenance of our mouse strains, the Flow Cytometry Facility, the Advanced Sequencing Facility, and the Histopathology Facility.

## Disclosure and competing interest statement

The authors have no competing interests.

---

**Expanded View 1-.**
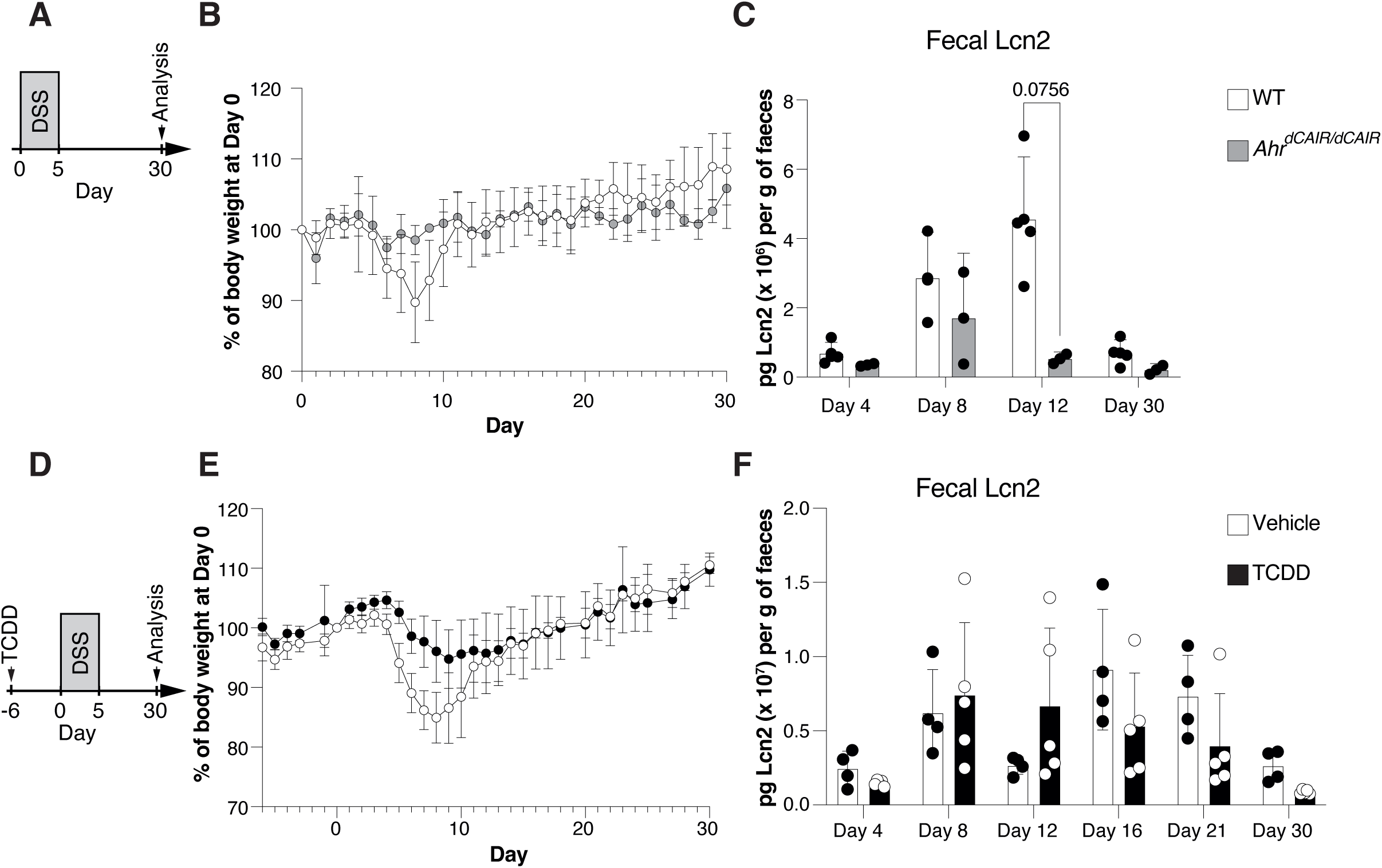
Prolonged AHR activation reduces DSS-induced inflammation. (**A, D**) Experimental scheme, (**B, E**) body weight curves and (**C, F**) Lcn2 levels in faecal samples at the indicated timepoints in (**A-C**) Ahr^CAIR/CAIR^ and WT mice (WT, n=4; *Ahr^dCAIR/dCAIR^*, n=2) and (**D-F**) TCDD and vehicle-treated mice (Vehicle, n=4, TCDD, n=5). Each dot represents the mean + SD (B, E) and bars represent the mean + SD (C, F). Two-way ANOVA with Šídák’s multiple comparisons test (C, F).

**Expanded View 2-.**
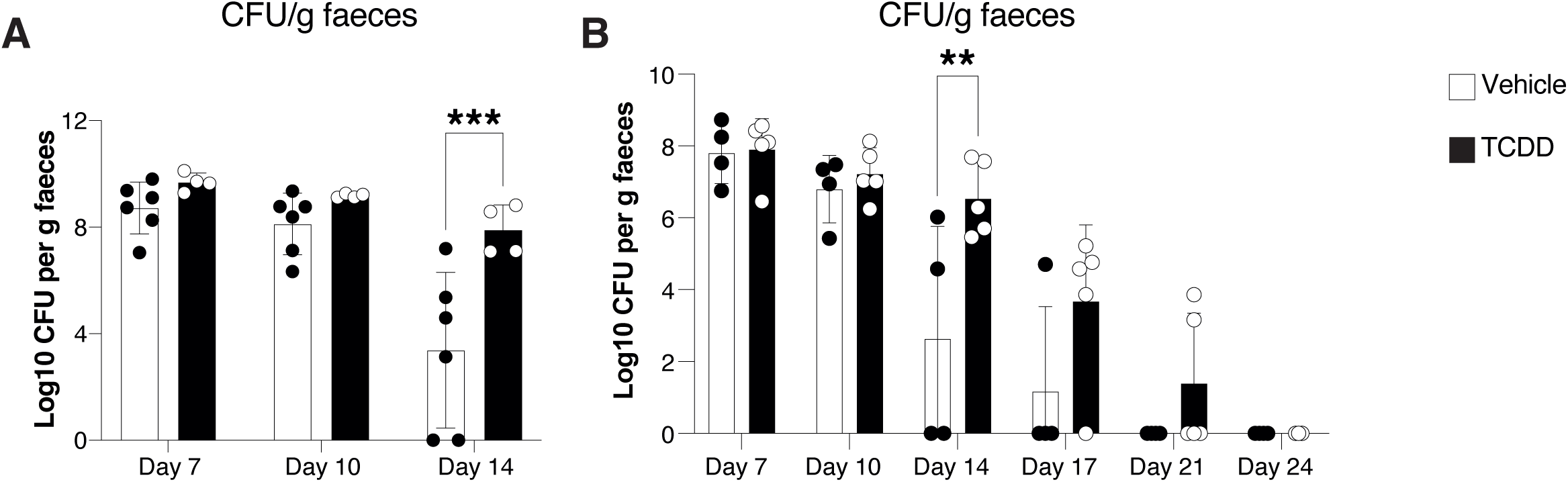
Delayed clearance of *C. rodentium* in mice exposed to TCDD. **(A-B)** *C. rodentium* burden in faecal pellets of female mice infected with *C. rodentium* 6 days after an oral dose of TCDD or vehicle. Data are from 1 experiment with n=4-6 per group. (**B**) *C. rodentium* burden in faecal pellets of mice infected with *C. rodentium* 6 days after an oral dose of 10 μg/kg TCDD or vehicle before infection. Data are representative of 2 experiments with n=4-5 mice per group. Each dot corresponds to one mouse and bars represent the mean + SD. Two-way ANOVA with Šídák’s multiple comparisons test (A, B). *p<0.05, **p<0.01, ***p<0.001.

**Expanded View 3-.**
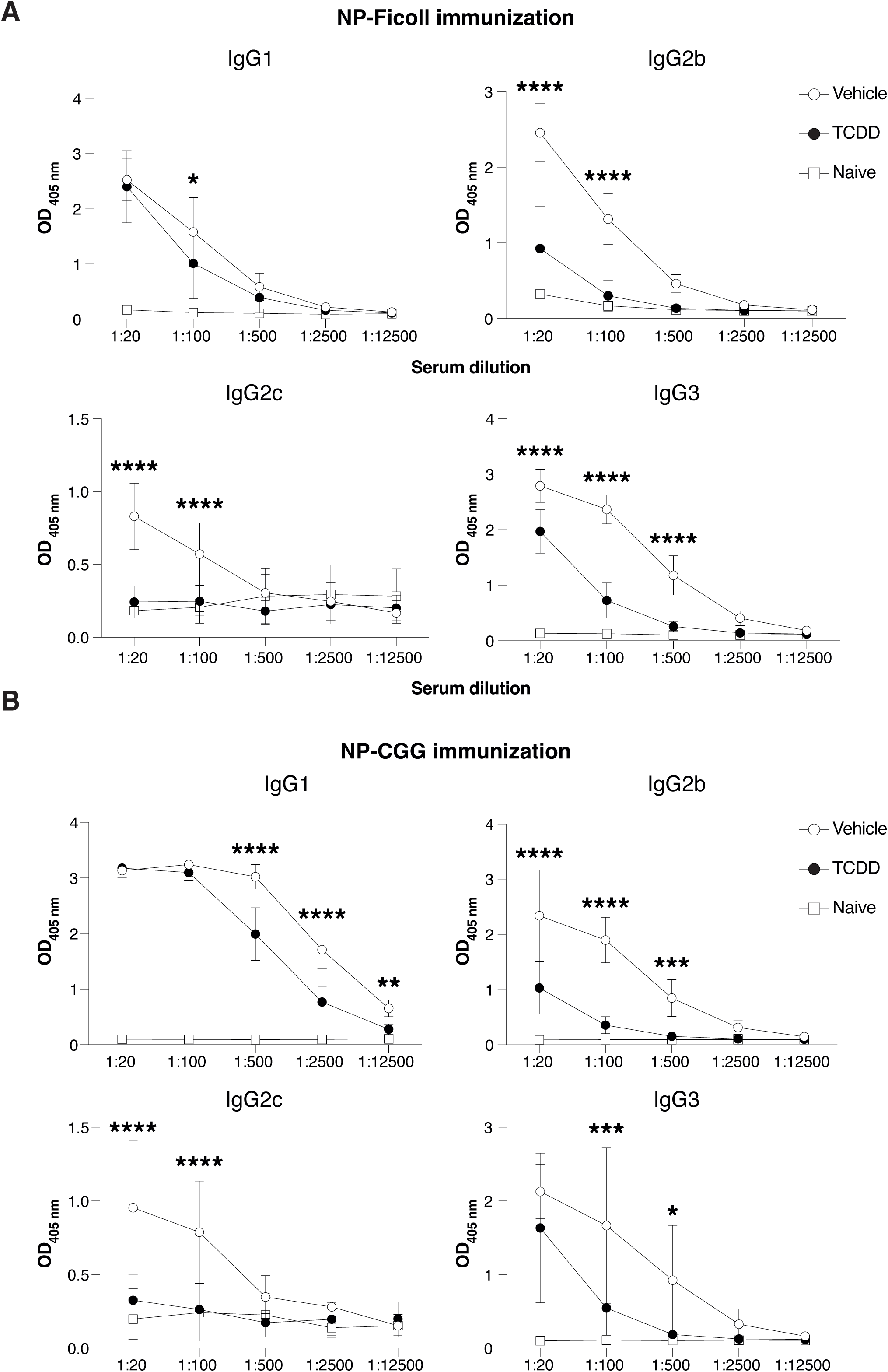
TCDD suppresses production of NP-specific IgG antibodies. (**A**, **B**) Levels of NP-Ficoll and NP-CGG-specific IgG subclasses at the indicated sera dilutions 14 days after immunization with 10 μg NP-Ficoll or NP-CGG, respectively. Mice received 10 μg/kg TCDD or vehicle 6 days before immunization. Data are representative of 2 experiments (Vehicle, n=10; TCDD, n=10, Naïve =2). Each dot represents the mean of 10 mice + SD. Two-way ANOVA with Tukey’s multiple comparisons test (A-B). *p<0.05, **p<0.01, ***p<0.001, ****p<0.0001.

**Expanded View 4-.**
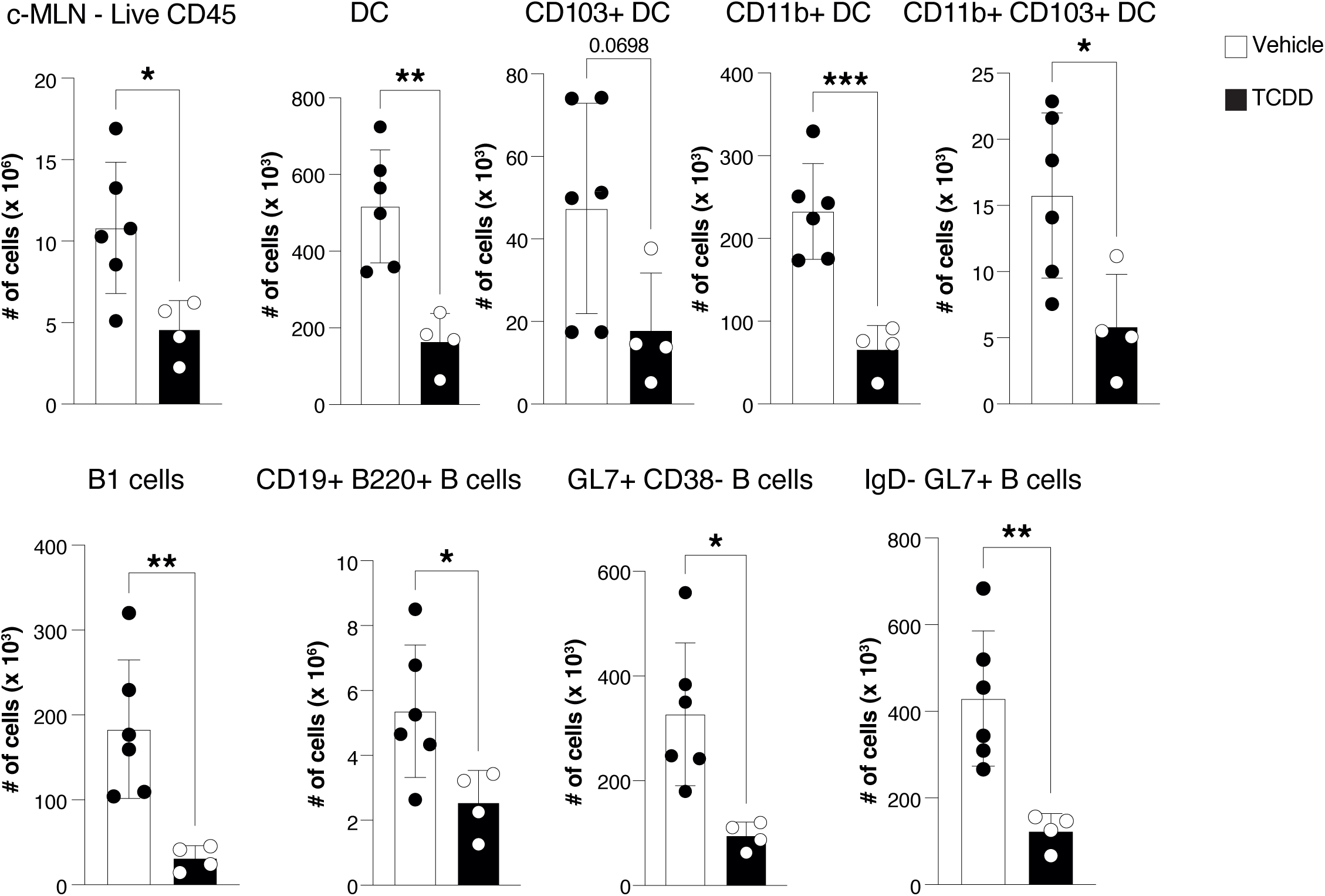
TCDD affects immune cell composition in the c-MLN. Cell numbers of total immune cells, and dendritic cell and B cell populations in the c-MLN of mice infected with *C. rodentium* 6 days after receiving 10 μg/kg TCDD or vehicle. Data analysed by flow cytometry representing 1 experiment (vehicle, n=6; TCDD, n=4). Each dot represents one mouse, and the bars show the mean + SD. Student’s t-test. *p<0.05, **p<0.01, ***p<0.001.

**Expanded View 5-.**
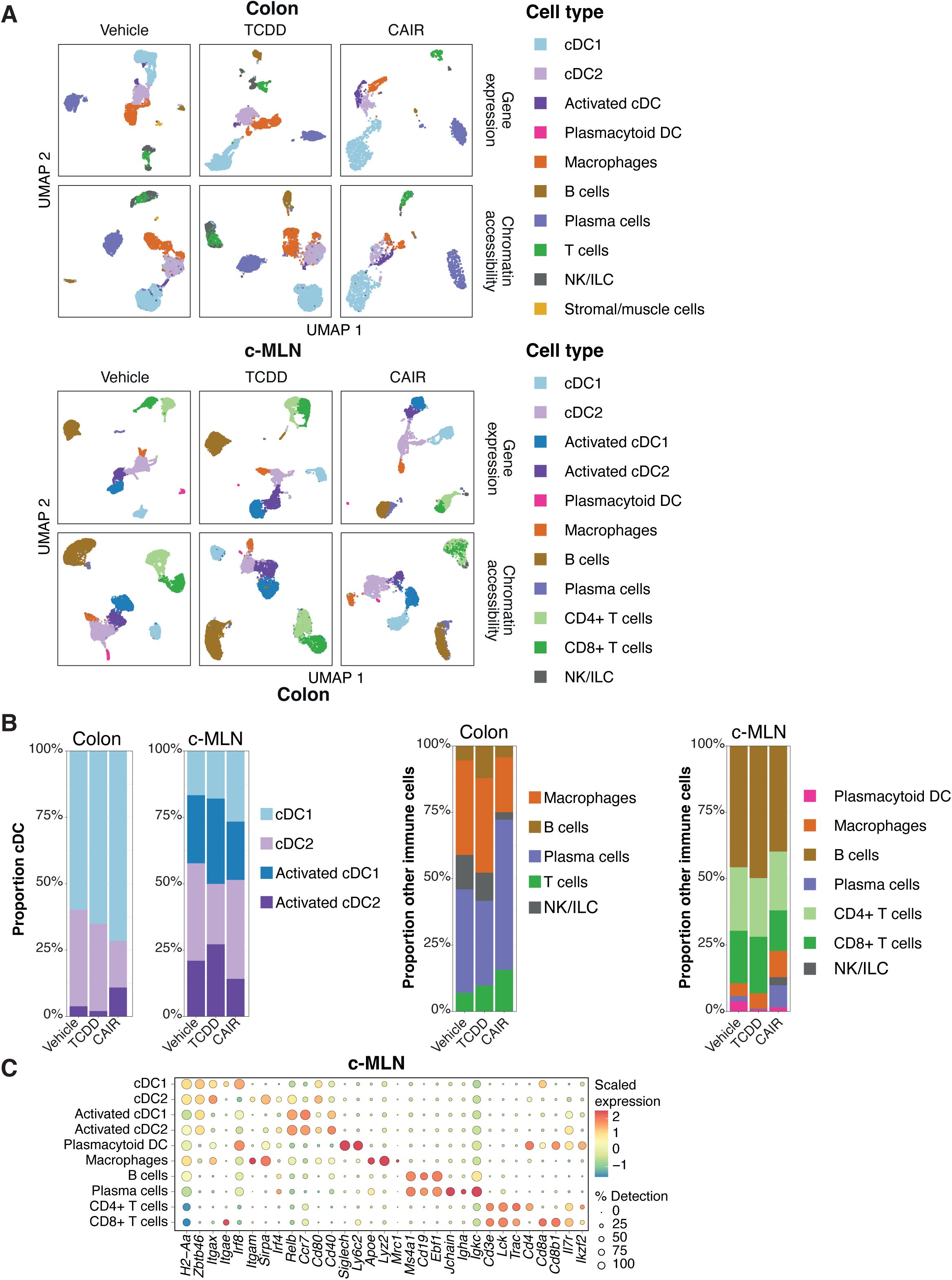
Cell types identified in the single cell RNAseq and ATACseq datasets. **(A)** Uniform manifold approximation and projection (UMAP) using scRNA-seq or chromatin accessibility. Cell types were assigned using the RNA-seq assay and visualised on both RNA- and ATAC-seq projections. **(B)** Proportion of dendritic cell subpopulations and of all other immune cells. **(C)** Dot plots adapted from Seurat DotPlot showing the putative marker genes across cell types defined in the vehicle sample in the MLN. The size of the dot represents the percentage of cells within a cell type expressing the marker gene, while colour represents the average scaled expression level of the marker gene in the cell type.

